# A catalogue of resistance gene homologs and a chromosome-scale reference sequence support resistance gene mapping in winter wheat

**DOI:** 10.1101/2022.01.26.477808

**Authors:** Sandip M. Kale, Albert W. Schulthess, Sudharsan Padmarasu, Philipp H. G. Boeven, Johannes Schacht, Axel Himmelbach, Burkhard Steuernagel, Brande B. H. Wulff, Jochen C. Reif, Nils Stein, Martin Mascher

**Author notes:** These authors contributed equally (Sandip M. Kale, Albert W. Schulthess). Correspondence should be addressed to Nils Stein or Martin Mascher.

## Abstract

A resistance gene atlas is an integral component of the breeder’s arsenal in the fight against evolving pathogens. Thanks to high-throughput sequencing, catalogues of resistance genes can be assembled even in crop species with large and polyploid genomes. Here, we report on capture sequencing and assembly of resistance gene homologs in a diversity panel of 907 winter wheat genotypes comprising *ex situ* genebank accessions and current elite cultivars. In addition, we use accurate long-read sequencing and chromosome conformation capture sequencing to construct a chromosome-scale genome sequence assembly of cv. Attraktion, an elite variety representative of European winter wheat. We illustrate the value of our resource for breeders and geneticists by (i) comparing the resistance gene complements in plant genetic resources and elite varieties and (ii) conducting genome-wide associations scans (GWAS) for the fungal diseases yellow rust and leaf rust using reference-based and reference-free GWAS approaches. The gene content under GWAS peaks was scrutinized in the assembly of cv. Attraktion.

## Introduction

Maintaining plant health in the face of evolving pathogen populations is a perennial goal of breeders. Key to this endeavor is the discovery and deployment of disease resistance (R) genes. Hafeez et al. (2021) put forward the concept of an R gene atlas and illustrated its potential for crop improvement in one of our most widely grown crops, wheat. An important component of populating the wheat R gene atlas is genotyping diversity panels, or more broadly, knowledge of as large a fraction of the resistance gene complement of as many genotypes as possible. One approach to this aim, resistance gene enrichment sequencing [RenSeq, Jupe et al. (2013)], was developed with large-crop genomes in mind. To reduce genomic complexity of sequencing libraries, and hence the required sequence effort, capture probes are designed to target R gene homologs from the nucleotide-binding and leucine-rich repeat (NB-LRR) family, or more generally the family of NB-LRR-related genes (NLRs, Ting et al. (2008)). In its original implementation, RenSeq was combined with short-read sequencing on the Illumina platform (Jupe et al., 2013). A combination of RenSeq with long-read sequencing has been used to assemble the full complement of R genes in the model plant *Arabidopsis thaliana* and analyze their evolutionary dynamics (Van de Weyer et al., 2019).

RenSeq data for diversity panels in combination with matching phenotype data has been used for genome-wide associations scans (GWAS) to find genetic markers associated with disease resistance (Arora et al., 2019). In the best case, this method, termed AgRenSeq, can zoom in on individual candidate genes. However, the limits of association mapping such as population structure (Yu et al., 2006) and sensitivity to the genetic architecture of the trait under study (Lopez-Arboleda et al., 2021) also apply to AgRenSeq. Recently, Gaurav et al. (2021) reported the use of whole-genome shotgun sequencing for association mapping of disease resistance in the wheat diploid progenitor *Aegilops tauschii*. An advantage of WGS over RenSeq is its ability to access also non-NLR resistance genes; a potential drawback is the inability to assemble full-length genes from low to medium-coverage (3x-10x) short-read data.

Independent of choice of sequencing strategy, a potential impediment to GWAS as well as a crucial aspect of R gene evolution is structural variation (SV). R genes are subject to ubiquitous presence-absence and copy-number variations (Michelmore and Meyers, 1998; Van de Weyer et al., 2019). Reference-free GWAS approaches have shown that the presence of peaks can be influenced by the choice of reference sequence (Voichek and Weigel, 2020). In principle, the best resource for studying intra-species NLR diversity are high-quality genome assemblies for a representative diversity panel comprising hundreds of accessions, i.e. a pan-genome. Constructing pan-genome infrastructures for all major crops has recently turned from a moon shot into a realistic mid-term research goal (Della Coletta et al., 2021). But the wheat pan-genome is not there yet: chromosome-scale reference genome sequences for ten wheat varieties, most of them recent elite cultivars, have recently been released (Walkowiak et al., 2020), but this small panel is not comprehensive enough to underpin a species-wide resistance gene inventory.

In the present manuscript, we report on a contribution to the wheat R gene atlas. We constructed an R gene inventory for a diversity panel of winter wheat, the predominant type of wheat in Europe. A chromosome-scale reference genome sequence was constructed for one representative winter wheat cultivar. To illustrate the value of this resource for the wheat genetics and breeding community, we (i) compare patterns of R gene diversity between plant genetic resources (PGR) and elite cultivars; (ii) conduct GWAS for the fungal diseases yellow rust and leaf rust; and (iii) analyze structural variants in close proximity to significantly associated markers.

## Results

### R gene capture in a winter wheat diversity panel

We conducted RenSeq for a panel of 907 winter wheat genotypes (**Figure 1, Table S1**) and the reference genotypes Chinese Spring (The International Wheat Genome Sequencing Consortium (IWGSC), 2018) and Julius (Walkowiak et al., 2020). Of these, 779 are part of a previously described core set enriched for disease resistant genotypes (Schulthess et al., 2021) comprising 587 PGRs and 192 European elite cultivars. The remaining 128 genotypes are recent German elite breeding lines. We used the Triticeae RenSeq Baits V3 (Tv3) probe set comprising 217,827 oligonucleotide baits (Zhang et al., 2021). Alignment of this bait set to the Chinese Spring reference genome (RefSeq v1.0, The International Wheat Genome Sequencing Consortium (IWGSC) (2018)) indicated that 18 Mb of annotated NBS-LRR gene sequence are targeted. On average, sequences originating from the predicted target were enriched 220-fold.

**Figure 1:**
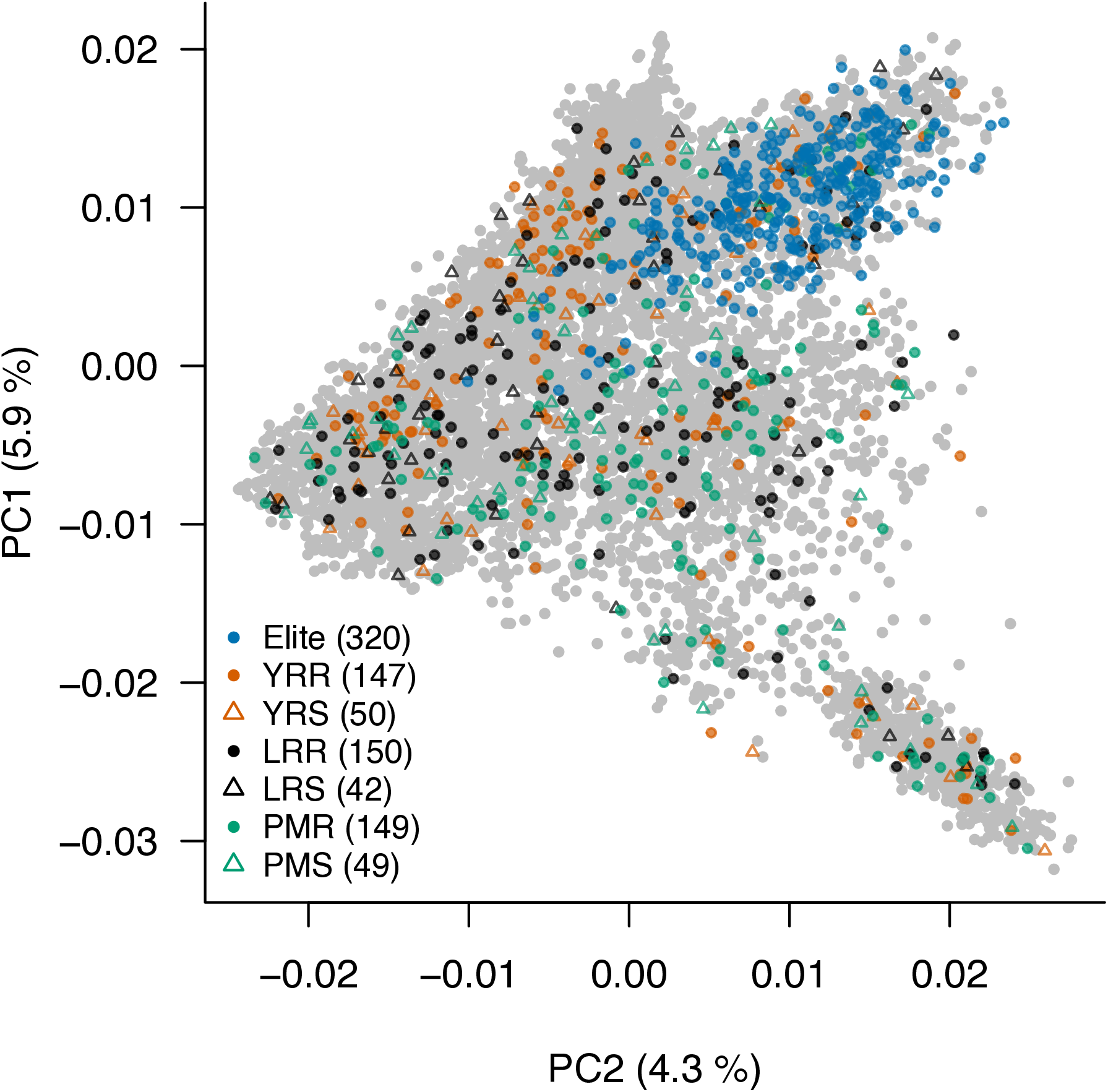
Genetic diversity of genotypes selected for RenSeq. The 907 accessions selected for the RenSeq analysis were projected onto the molecular diversity space of the winter wheat collection of the IPK genebank portrayed by the first two principal components (PCs) from a PC analysis on genome-wide SNP markers (Schulthess et al. 2021). Among RenSeq characterizations, 192 European elite cultivars and 128 German elite breeding lines represent the diversity already handled by European breeding. The remaining fraction is composed of 587 plant genetic resources (PGRs) samples from the IPK genebank which was enriched for disease resistant genotypes with minimized population structure (Schulthess et al. 2021). According to selection, PGRs are classified as yellow rust resistant [YRR] or susceptible [YRS], leaf rust resistant [LRR] or susceptible [LRS] and powdery mildew resistant [PMR] or susceptible [PMS].

RenSeq reads of individual genotypes were assembled *de novo*, yielding 67,731 to 4,583,579 contigs per accession (**Table S2**). Of these, 417 to 2,304 per accession (mean: 1,690) contained full-length NLRs. Coiled-coil NLRs were the most abundant class of NLRs (**Figure 2**). Almost all (1,911/1,937) NLRs assembled from the Chinese Spring RenSeq data were aligned to RefSeq v1.0 with 95 % alignment coverage and 95 % alignment identity. A total of 1,486 (77 %) Chinese Spring *de novo* assembled NLRs overlapped with RefSeq v1.0 gene models, indicating the absence of resistance gene homologs in the reference annotation, possibly because of lack of expression or pseudogenization. In other genotypes, on average 77 % of assembled NLRs were mapped to RefSeq v1.0, consistent with pervasive presence-absence variation (PAV) in resistance genes.

**Figure 2:**
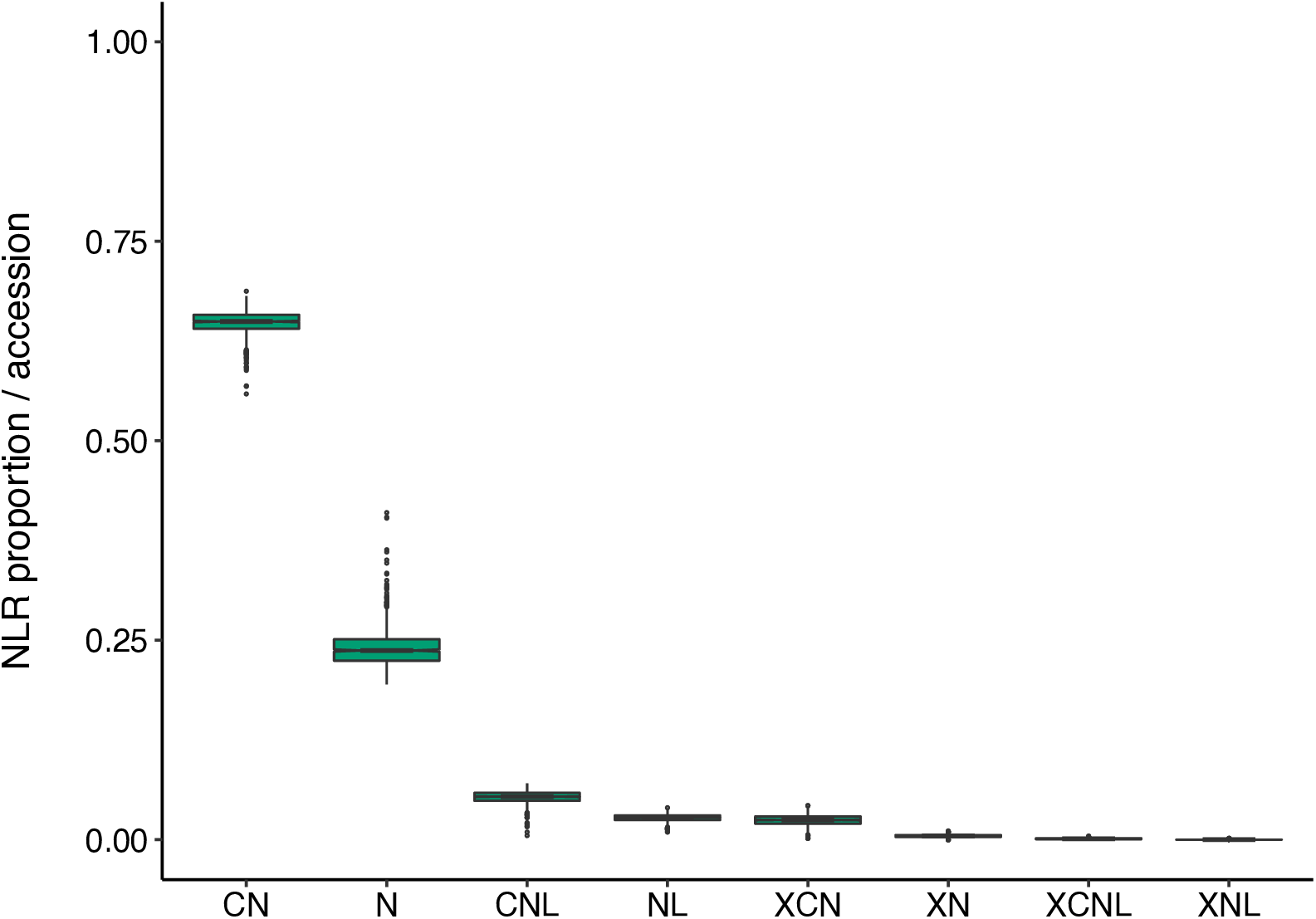
Proportion of NLRs of different classes in individual RenSeq assemblies. CN: Colied-coil-NBS; N: NBS; CNL: Colied-coil-NBS-LRR; NL: NBS LRR; XCN: integrated domain (ID)-Colied-coil-NBS; XCNL: ID-Colied-coil-NBS LRR; XNL: ID-NBS-LRR.

### Diversity of NLRs in winter wheat genepools

To understand the extent of PAV in NLRs in our panel, we performed similarity-based clustering of the assembled NLR from all genotypes. A total of 1,469,694 (96 %) of NLRs were clustered in 39,073 orthogroups, the remainder were singletons without close matches to other NLRs. Fewer than 1 % of orthogroups contained two or more NLRs from the same accession, pointing to a potential collapse of highly similar, recently duplicated NLRs. Most (> 85 %) of orthogroups had NLRs from at least 20 different accessions. However, very few orthogroups (472, 1.21%) had members from more than 500 accessions (**Figure S1**). This is at odds with patterns of NLR diversity in the model plant *Arabidopsis thaliana* (Van de Weyer et al., 2019), where the “core-NLRome” comprising genes present in almost all genotypes is substantial. The likely explanation is random sequence dropout due to competition between capture probes and/or low sequencing depth. For example, at a 1 % dropout rate (i.e., a 99 % change of being captured and sequenced at sufficient depth), a gene present in 900 genotypes has a negligible chance (0.99^900^ = 0.01 %) of being present in all their assemblies. A saturation analysis indicated that a near-complete set of NLR orthogroups assembled in the whole panel can be captured with a rather small number of accessions: 95 % of orthogroups were captured with only 70 genotypes selected at random from the universe of 907 accessions (**Figure 3**). Because of random dropout, these figures are likely overestimates, i.e. an even smaller panel may suffice to reach the 95 % threshold. Still, the analysis of orthogroups allows comparison of relative diversity between gene pools. When considering PGR and elite accessions separately, near-saturation can be achieved with 80 and 150 accessions, respectively, indicating that, not unexpectedly, NLR diversity is higher in PGRs. However, elite lines were more resistant against yellow rust compared to PGRs and contained a higher number of NLRs that preferentially occur in resistant genotypes, supporting the notion that breeder’s efforts to stack resistance genes have been successful (**Figure 4a**). A potential caveat, though, is that NLRs private to elite varieties were localized to regions previously reported as harboring alien introgressions. By contrast, PGR-specific NLRs were distributed more uniformly across the chromosomes (**Figure 4b**). Hence, most NLRs occurring in highly resistant elite varieties may not confer resistance on their own, but have only hitchhiked along one or a few functional resistance genes targeted by breeders.

**Figure 3:**
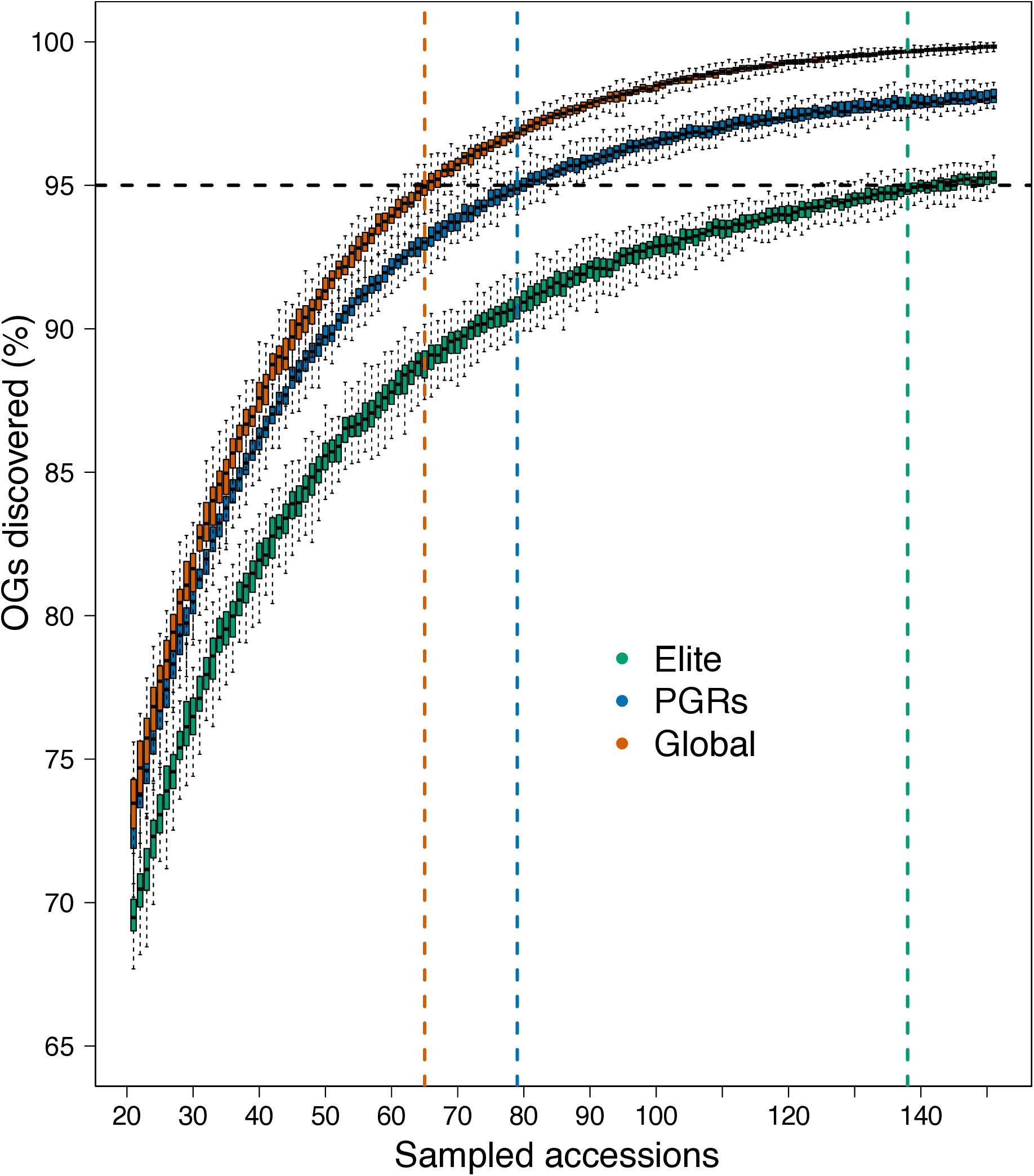
Saturation analysis. Fraction of NLR orthogroups recovered from randomly drawn subsets of genotypes. Subsets were selected from the entire population as well as elite varieties and PG Rs. Sampling was repeated 100 times for subsets of increasing size. Colored vertical lines indicate the number of accessions required to achieve 95 % representation of the NLR universe.

**Figure 4:**
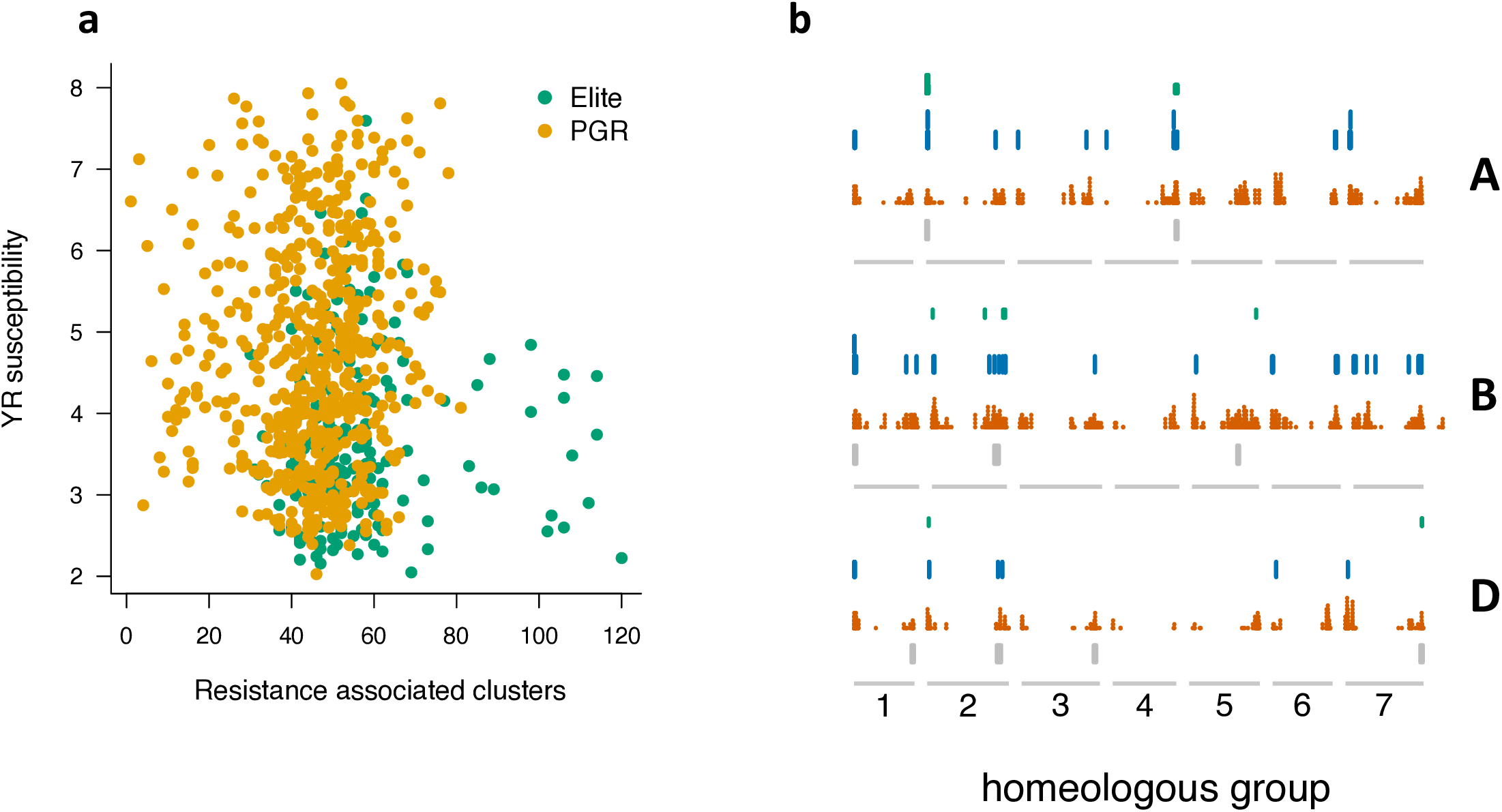
Enrichment of resistance-associated NLR clusters in elite lines. **(a)** The number of resistance-associated clusters from an accession is plotted against its yellow rust susceptibility score. Elite lines with many resistance-associated clusters were less susceptible to yellow rust. **(b)** Genomic distribution of elite specific orthogroups (OGs, green), PGR-specific OGs (blue) and OGs present in both elite lines and PGRs (orange) in the three subgenomes of hexploid wheat (A, B, D). The grey boxes mark the positions of alien introgressions.

### A chromosome-scale assembly of cv. Attraktion

Several recent studies suggest that the choice of reference genome impacts the contextualization or even the very presence of GWAS peaks (Arora et al., 2019; Voichek and Weigel, 2020). It is likely that a better reference genome than the assembly of Chinese Spring – indeed a spring-sown landrace from China (Sears and Miller, 1985) – can be selected for mapping resistance genes in winter types. We chose cv. Attraktion because our prior analysis of shallow-coverage whole-genome shotgun data had shown that this cultivar carries large alien introgressions, some of them co-incident with GWAS peaks for resistance to yellow and leaf rust (Schulthess et al., 2021).

We sequenced the Attraktion genome to 22-fold coverage with HiFi reads with an average length of 17.8 kb of circular consensus reads. In addition, chromosome conformation capture sequencing (Hi-C) was performed, resulting in 994 million read pairs. Genome assembly following a previously described approach (Mascher et al., 2021; Sato et al., 2021) combining primary contig assembly with Hifiasm (Cheng et al., 2021) and pseudomolecule construction with the TRITEX pipeline (Monat et al., 2019) yielded a set of 1,553 contigs (14.25 Gb) assigned to chromosomal locations. A further 3,442 contigs (434 Mb) remained unplaced. A BUSCO analysis (Simao et al., 2015) indicated that 98.2 % of conserved single-copy genes were present in the assembly (**Table 1**). The inspection of Hi-C contact matrices and alignment to the Chinese Spring RefSeq v2.1 (Zhu et al., 2021b) supported the structural integrity of the pseudomolecules (**Figures S2, S3**).

**Table 1:** Statistics of the genome sequence assembly of cv. Attraktion.

|  |  |
| --- | --- |
| Assembly size | 14.7 Gb |
| Number of contigs | 4,953 |
| Contig N50 | 17.3 Mb |
| Contig N90 | 4.1 Mb |
| Pseudomolecule size | 14.3 Gb |
| Number of contigs in pseudomolecules | 1,553 |
| Complete BUSCOs | 1,584 (98.2%) |

Regions of high divergence between Attraktion and Chinese Spring indicative of the presence of alien introgressions were found on four chromosomes: 4A, 2B, 5B and 2D (**Figure 5**). Interestingly, a 55 Mb introgression on the long arm of chromosome 2B in Attraktion overlapped with a much larger 427 Mb introgression from *T. timopheevi* in LongReach Lancer. Attraktion and Lancer have the same haplotype in the overlapping region, pointing to shared ancestry. Most likely, breeders had decreased the size of this introgression in Attraktion in an attempt to reduce linkage drag.

**Figure 5:**
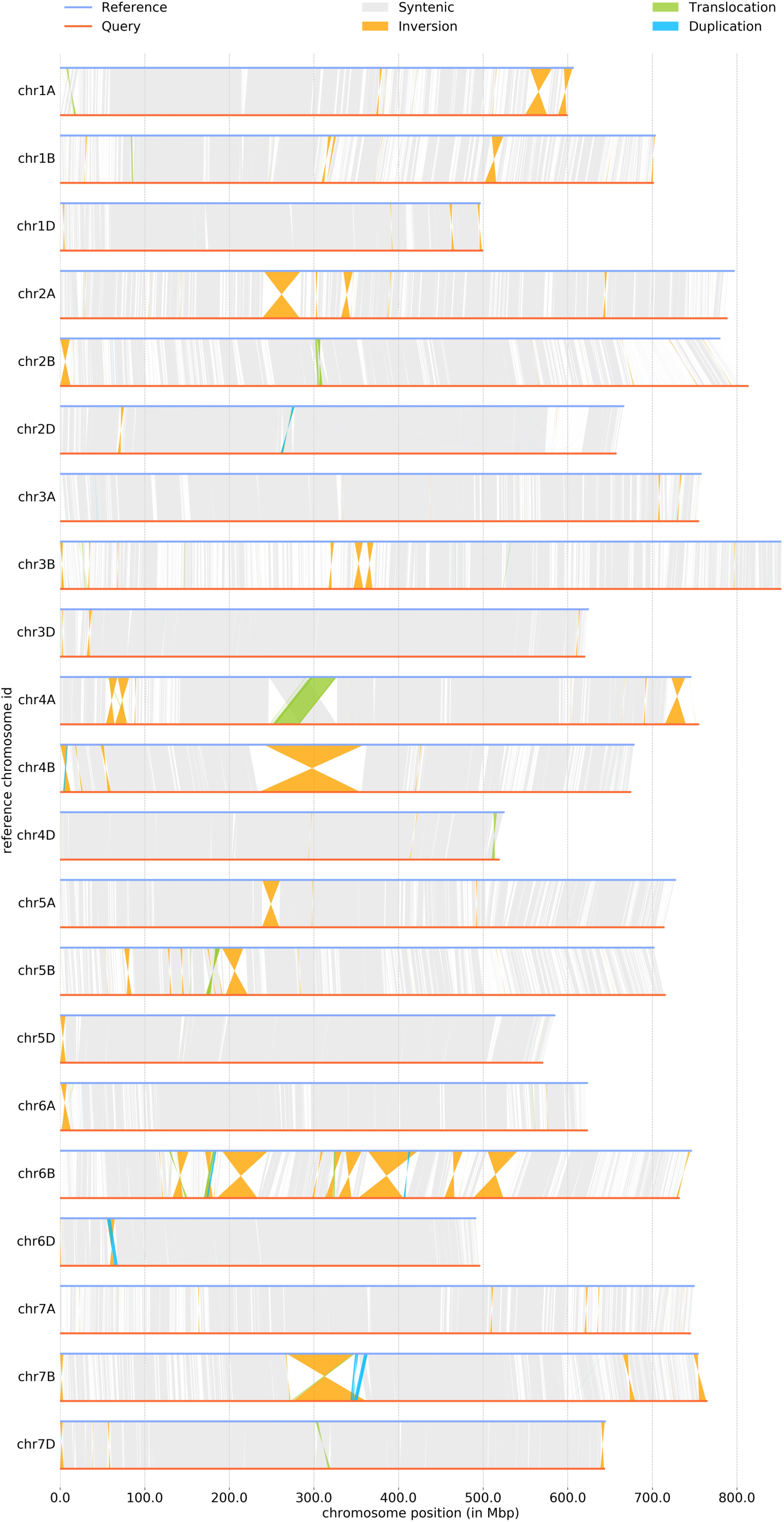
Structural variants detected by Synteny and Rearrangement Identifier (SyRI, Goel et al. 2019) between the genome assemblies of cv. Attraktion (reference) and Chinese spring RefSeq V2.1 (query).

### Different GWAS approaches identify a yellow rust resistance locus on chromosome 6A

To illustrate the value of our resource for genetic mapping of disease resistance, we conducted GWAS for yellow rust (*Puccinia striiformis* f.sp. *tritici*) resistance in our panel. The degree of yellow rust infection was scored in multi-environment field trials, relying on natural and artificial infection. Details are described elsewhere (Schulthess et al., 2021). We followed three different approaches to obtain matrices of bi-allelic markers for use in GWAS. First, we aligned RenSeq reads to a reference genome sequence assembly, called single-nucleotide polymorphisms (SNPs), and used genotype calls at SNP sites as markers. This is the most commonly applied approach for marker discovery, which, however, can capture structural variants only if they are in close linkage disequilibrium (LD) with SNPs. Second, we conducted kmerGWAS (Voichek and Weigel, 2020) which queries the presence-absence state of short oligonucleotides of a fixed length (*k*-mers, *k*=31) as proxies for structural variants. Third, we used SNP sites discovered from the alignment to the reference assembly, but instead of allelic status, we used presence-absence states of genotype calls as markers, similar to what Gabur et al. (2018) did with SNP chip data of rapeseed. We refer to this method as paGWAS. Two different reference genome sequences, Chinese Spring RefSeq V2.1 and our Attraktion assembly, were used to position markers. Note that kmerGWAS is a reference-free approach; associated *k*-mers were aligned *post hoc* to the genome assemblies to place them.

Manhattan plots for all three methods and the two references are shown in **Figure 6**. The most prominent feature is a peak on the long arm of chromosome 6A, for which significantly associated markers were reported in GWAS scans. However, it is less prominent in paGWAS and kmerGWAS against the Chinese Spring reference, possibly reflecting the absence of the resistant haplotype in that genotype. SNP GWAS against Chinese Spring did result in a pronounced peak, likely because of SNPs in linkage disequilibrium with the causal variant. Further peaks were observed on other chromosomes, but were not common between all methods.

**Figure 6:**
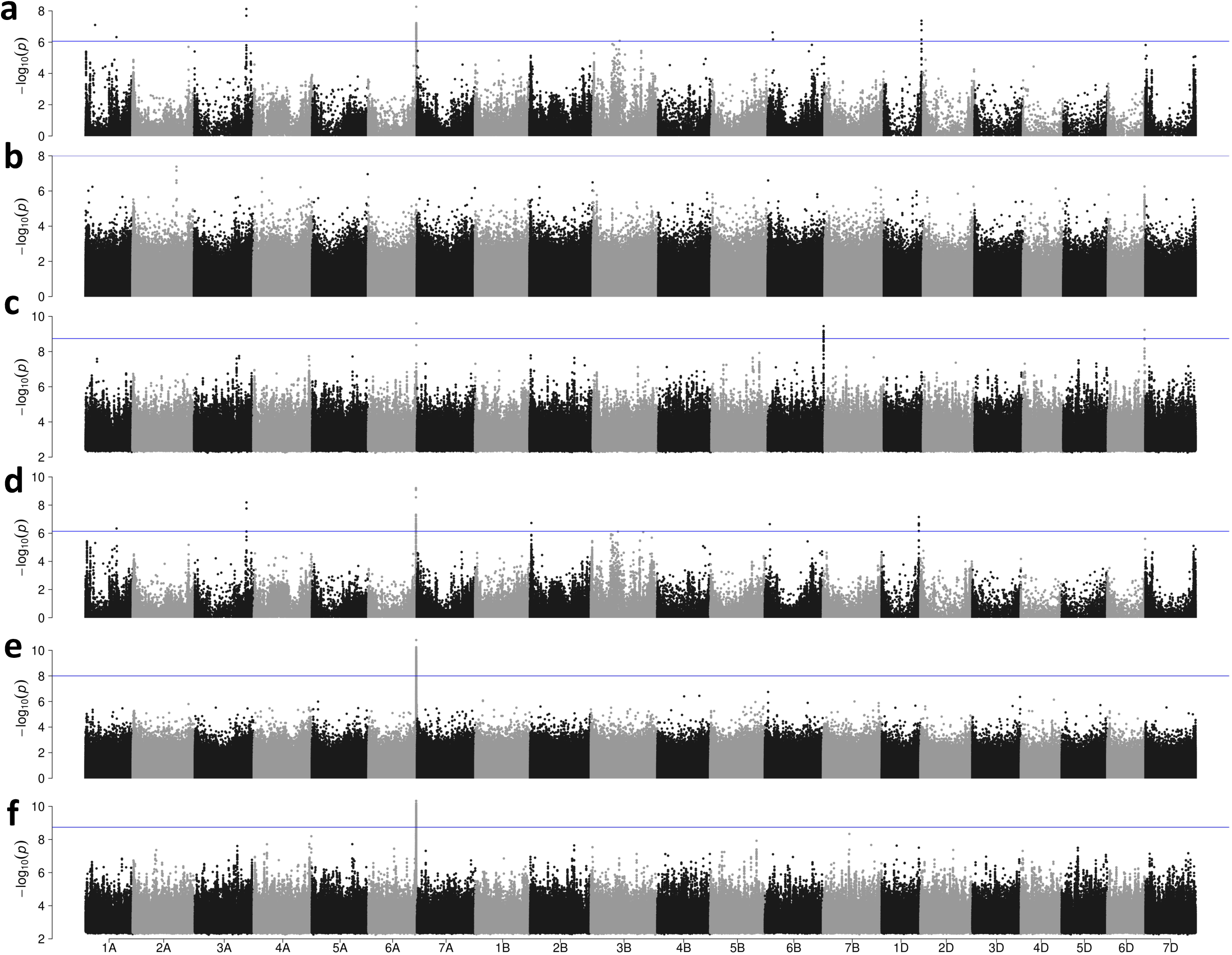
Association scans for yellow rust resistance using different marker systems. **(a)** SNPs identified relative to Chinese Spring RefSeq Vl.O. **(b)** presence-absence GWAS using SNPs identified relative to Chinese Spring RefSeq Vl.O and scored as presence-absence markers. **(c)** *k*-mers mapped against Chinese Spring RefSeq Vl.O. Panels **(d), (e)** and **(f)** show the results of SNP-based, presence-absence and *k*-mer based GWAS when the reference sequence of cv. Attraktion was used for SNP identification or *k*-mer mapping. The blue horizontal lines indicate the threshold above which associations are statistically significant.

To the best of our knowledge, a resistance gene against yellow rust has not been reported on chromosome 6A in the region pinpointed by our GWAS. We scrutinized the region under the peak in the genome assembly of cv. Attraktion, which scored highly in our resistance trials and carried the resistant haplotype at the 6A peak. The significantly associated markers spanned an interval of 946 kb in the Attraktion genome (**Figure 7a**), containing 121 gene models annotated *ab initio* (Stanke et al., 2006), many which are actually derived from transposable elements. Seven of these genes were NLRs. One of them, spanning a 4.9 kb at around sequence coordinate 612.5 Mb on the 6A pseudomolecule of Attraktion, was identical to the representative contig of the orthogroup “cluster49707” identified from the RenSeq *de novo* assemblies. This representative contig harbored 46.9 % of the significantly associated 31-mers. Among the 10 genome sequence assemblies reported by Walkowiak et al. (2020), that of SY Mattis had the same haplotype as Attraktion (**Figure 7a**). The other nine genomes as well as the Chinese Spring reference lacked several genes present in the Attraktion haplotype, including cluster49707.

**Figure 7:**
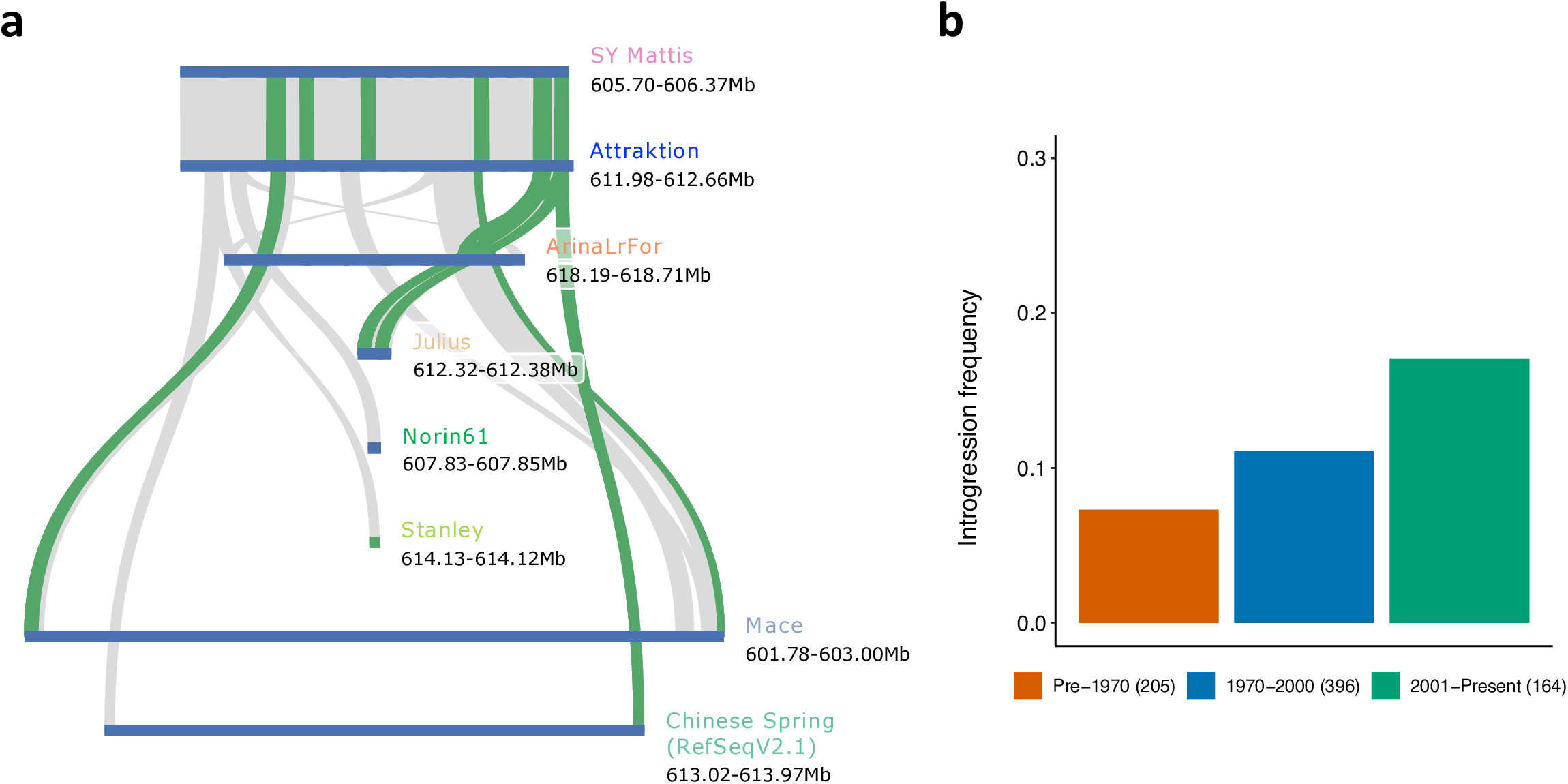
Tracing the history of a novel yellow rust resistance locus on chromosome 6A. **(a)** Gene-based colinearity analysis of the yellow rust resistance locus identified on chromosome 6A in the Attraktion assembly with reference assemblies from the wheat pan-genome (Walkowiak et al. 2020). **(b)** Frequency of resistant haplotypes in accessions from different time period. The numbers in parentheses indicate size of each group.

Interestingly, no significant marker-trait associations were detected in the peak region when only PGRs were included in the association scan (**Figure S4**). The high synteny between Chinese Spring and Attraktion in the vicinity of the peak rules out the presence of an alien introgression. We speculate that the resistant haplotype may have segregated at low frequency – below the detection threshold of GWAS – in landraces and old varieties. Recent shifts in European pathogen populations (Hovmøller et al., 2016) may have favored the resistant haplotype in European winter wheats (**Figure 7b**). Future work should focus on the identification and validation of the causal gene conferring yellow rust resistance and on singling out the yellow rust isolates that is recognizes.

### GWAS for leaf rust detects known and novel loci

The second trait for which we did GWAS is leaf rust (*Puccinia triticina* f.sp. *tritici*) resistance. The degree of natural infection with leaf rust was scored under field conditions (**Tables S3 and S4**). SNP, paGWAS and kmerGWAS gave partially overlapping results (**Figure 8a, Supplementary Figure S5**). Common to all approaches was a peak towards the distal end of the long arm of chromosome 4A. This association had been reported before by Liu et al. (2020a), who analyzed 133 genotypes and 1,574 of their hybrid offspring by exome sequencing. The GWAS peak was co-located with a 26 Mb region of high sequence divergence between Chinese Spring and Attraktion (**Figure 5**), which we attribute to an alien introgression, possibly originating from *T. dicoccoides* (Przewieslik-Allen et al., 2021). Because of suppressed recombination, the introgression is inherited as one large linkage block (**Figure 8b**).

**Figure 8:**
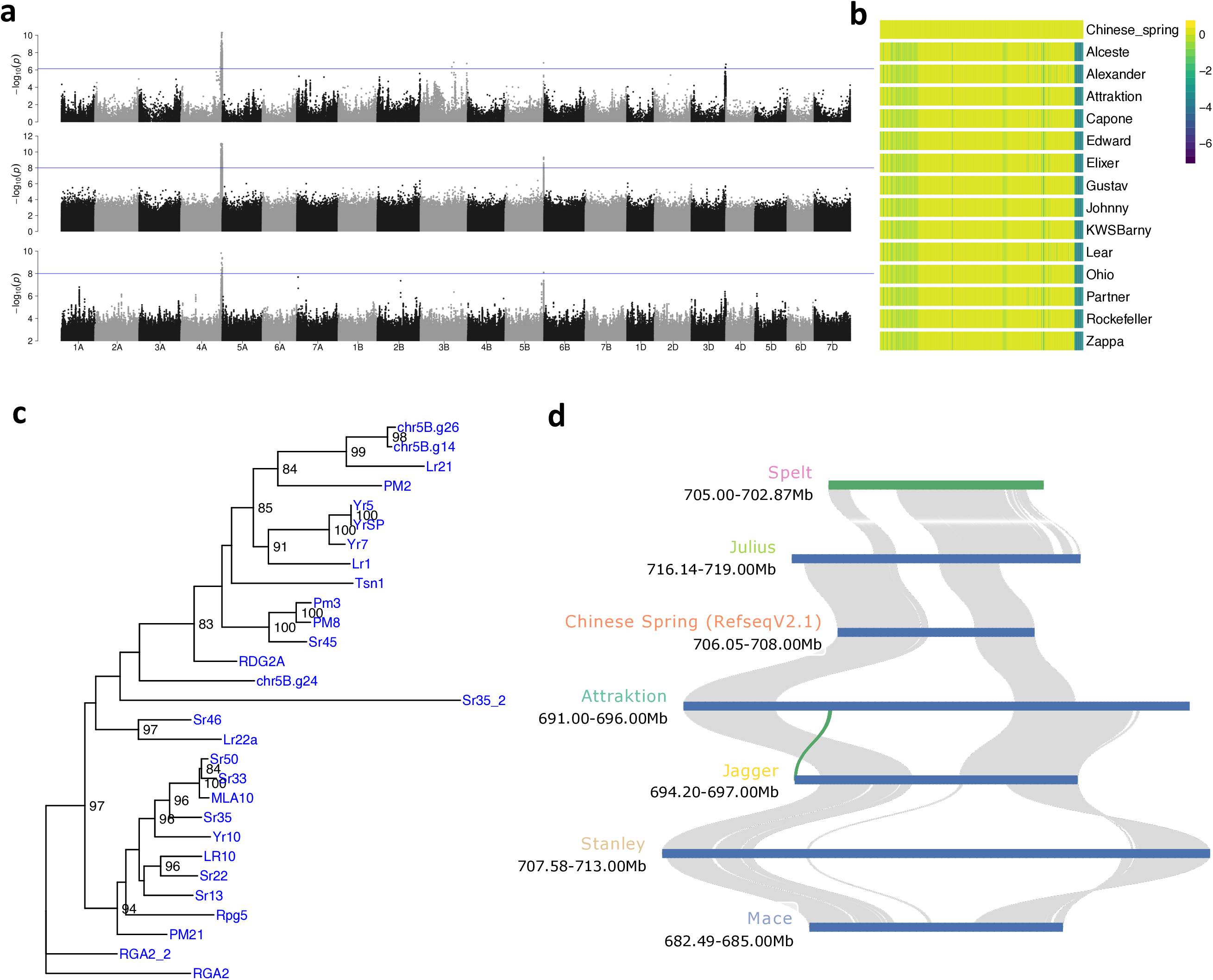
Identification of a leaf rust resistance locus. **(a)** Manhattan plots showing GWAS results for leaf rust resistance based on SNPs (top row), SNPs scored as presence-absence markers (middle row) and *k*-mer markers. SNPs and *k*-mers were anchored to the reference sequence assembly of cv. Attraktion. Two regions at the distal ends of the long arms of chromosomes 4A and SB were associated with leaf rust resistance. **(b)** Normalized read depth in 500 kb bins along chromosome 4A of Chinese Spring RefSeq Vl.O for representative elite varieties. **(c)** Phylogenetic tree constructed with NBS-LRR genes from the SB locus together with cloned resistance genes of wheat. Two genes from the locus are highly homologous to *Lr21*. **(d)** Gene based-colinearity analysis of the Attraktion haplotype at the SB locus with assemblies from the wheat pan-genome (Walkowiak et al. 2020), none of which carry the Attraktion haplotype.

Significant associations on the long arm of chromosome 5B were detected by multiple GWAS approaches. Cultivar Attraktion had high resistance scores and had a haplotype associated with resistance in the peak region, which extended from about 692.4 Mb to 694.1 Mb in the Attraktion genome assembly, spanning 235 gene models. Of these, thirteen were NLRs. Significantly associated *k*-mers mapped to 15 orthogroups of NLRs *de novo* assembled from our RenSeq data, which corresponded to three gene models in the Attraktion assembly. One of them was highly similar to the cloned *Lr21* resistance gene (Huang et al., 2003). Interestingly, none of the wheat pangenome assemblies (Walkowiak et al., 2020) harbored this gene, illustrating the need for bespoke genome sequence assemblies to capture the gene content of resistance gene clusters. Further work is needed to prove the causal link between resistance to leaf rust and presence of the *Lr21*-related gene or other NLRs under the GWAS peak.

## Discussion

We have reported RenSeq assemblies as a component of the burgeoning wheat R gene atlas. Due to short-comings of our short-read capture sequencing approach, we were unable to construct a comprehensive NLR-ome as was done with long-reads in *A. thaliana* (Van de Weyer et al., 2019). However, our data set did reveal a contrasting repertoire of R gene homologs in elite varieties and PGRs, documenting breeders’ efforts at enhancing genetic resistance by selecting haplotypes bearing R genes.

The assembly of one recent elite cultivar, Attraktion, proved instrumental in the analysis of the gene content surrounding two GWAS peaks on chromosomes 6A and 5B for yellow rust and leaf rust, respectively. But no single genotype can capture all resistance genes and sequence assemblies of other genotypes will be required to zoom in on candidate genes against other diseases or other isolates of yellow rust. Fortunately, the cost for wheat whole-genome assembly has decreased substantially in recent years. The Attraktion assembly was completed within three months after selection of that genotype and cost approximately EUR 40,000. This shows that whole-genome assembly still entails large expenses, which, however, may constitute a worthwhile investment if candidate genes cannot be pinned down by cheaper alternatives. As long as capture and selective sequencing of large (> 500 kb) genomic regions has not become routine (López-Girona et al., 2020), whole-genome assembly is a viable alternative even if only a single locus is of interest. Reference genome sequences of diversity panels large enough for GWAS (i.e. 100-1000 genotypes) would render both RenSeq and WGS superfluous: the comprehensiveness and context afforded by genome assembly cannot be matched by short-read approaches. However, the large size of the wheat genome (15 Gb) makes the assembly of hundreds or thousands of genotypes cost-prohibitive at the time of writing. Consequently, the resources reported in this article will retain their usefulness in the medium term.

The three different GWAS approaches we took, SNP GWAS, paGWAS and kmerGWAS, were only partially concordant, highlighting the potential benefits that may be reaped from an integration of the GWAS and pan-genomics toolkits. Computational frameworks to construct and analyze pan-genome graphs are under active development (Hickey et al., 2020; Li et al., 2020). Reduced-representation approaches focusing on the single-copy or repeat-depleted part of the genome have been applied in soybean (Liu et al., 2020b) and barley (Jayakodi et al., 2020). It is unlikely, however, that single-copy sequence can represent copy-number variation in rapidly evolving resistance genes. Future algorithmic work should focus on the graph-based representation of pan-genomes for complex plant genomics, graph-based read mapping and GWAS with multi-allelic structural variants captured in pangenome graphs.

## Methods

### DNA extraction, library preparation and sequencing for RenSeq

A total of 779 genotypes from a trait-customized core collection of winter wheat (Schulthess et al., 2021) along with 128 advanced elite lines were used for resistance gene enrichment sequencing (RenSeq). DNA was extracted from a single leaf of about 10 cm length harvested from a 10 days old seedling using the DNeasy 96 Plant Kit (QIAGEN, Hilden, Germany) as per the manufacturer’s instructions. DNA quality and quantity were determined using a 0.8% agarose gel and Qubit fluorometer (Life Technologies, CA, USA). The RenSeq libraries were prepared using the protocol of Steuernagel et al. (2017) with minor technical modifications. Briefly, 1 µg DNA from each genotype was fragmented to ~500 bp size using the Covaris S2 (Covaris, MA, USA). The fragmented DNA was purified using 0.6X AMPure® XP beads (Beckman Coulter, IN, USA) according to the manufacturer’s instructions. Paired end libraries for Illumina sequencing were constructed using NEBNext® Ultra™ II DNA Library Prep Kit for Illumina® (New England Biolabs Inc, MA, USA) as per the manufacturer’s instructions, except AMPure® XP beads were used for all the purification and size selection steps. For PCR amplification, 10 µl of adapter ligated DNA from each genotype was used along with 25 µl 2x KAPA HiFi HotStart ReadyMix (Kapa Biosystems, MA, USA), 1 µl Index and Universal PCR Primer and 13 µl water. The library from each genotype was indexed using Unique Dual Index Primer Pairs (NEBNext Multiplex Oligos for Illumina) in order to perform multiplexed sequencing.

The enrichment of NBS-LRR DNA fragments was achieved through hybridization of PCR amplified genomic DNA libraries prepared above with 200K Triticeae NLR bait libraries (Tv3, Zhang et al. (2021)) available at https://github.com/steuernb/MutantHunter/blob/master/Triticea_RenSeq_Baits_V3.fasta.gz. The libraries were quantified using Qubit fluorometer (Life Technologies, CA, USA) and average fragment size was determined using the 4200 Tape Station (Agilent Technologies, CA, USA). The libraries from eight genotypes were pooled in an equimolar manner and hybridized with the bait library (myBaits-11; Arbor Biosciences, Ann Arbor, MI, USA). The hybridization reaction was carried out at 65 °C for 18 hours and the hybridized fragments were captured using MyOne Streptavidin C1 magnetic beads (ThermoFisher Scientific). The hybridization and capture of NBS-LRR fragments was performed according to MYbaits v4.0 protocol. Finally, PCR amplification of captured fragments was carried out using 2x KAPA HiFi HotStart ReadyMix and standard Illumina P5 and P7 primers. Twelve capture libraries (96 genotypes) were pooled in equimolar amounts, quantified using qPCR and sequenced (paired-end, 2 × 250 cycles) on the NovaSeq 6000 (Illumina).

### DNA extraction, library preparation and sequencing for PacBio HiFi sequencing

High molecular weight (HMW) DNA for PacBio circular consensus sequencing (CCS) was prepared from 100 one-week old seedlings of cultivar Attraktion following the protocol from Dvorak et al. (1988). Briefly, nuclei were extracted from ground leaves in a sucrose-based homogenization buffer. The protein contamination was removed by proteinase-K treatment and phenol:chloroform extraction. The HMW DNA was then spooled out of solution during sodium acetate and ethanol precipitation. Size profile of the extracted DNA was checked using Femtopulse system genomic DNA 165 kb kit (Agilent Technologies, CA, USA). Eight HiFi SMRTbell® libraries were prepared using the SMRTbell™ Express Template Prep Kit 2.0 according to the manufacturer’s instructions (Pacific Biosciences protocol: PN 101-853-100 Version 03, January 2020). In short, the protocol involves fragmenting the HMW DNA to a mean fragment length of 20 kb using Megaruptor 3 (Diagenode), followed by DNA damage repair, end repair/A-tailing and adapter ligation. Linear DNA fragments were removed by nuclease treatment of the SMRTbell libraries. Size selection of libraries was carried out using the Sage ELF system and fractions with 15-20 kb mean insert sizes were used for sequencing. Polymerase/insert complex formation and clean up was performed using Sequel II™ binding kit 2.0 based on manufacturer’s instructions. Sequencing was performed on 16 8M SMART cells using sequencing chemistry V2.0 and with a 2-hours pre-extension and 30-hours movie time setting. CCS reads were obtained with PacBio CCS software (https://github.com/PacificBiosciences/ccs).

Chromosome conformation capture sequencing (Hi-C) libraries were prepared from 1.5 g of leaf material from one-week-old seedlings of cv. “Attraktion” as per the protocol of Padmarasu et al. (2019) with a few modifications. The modifications include the use of nuclei isolation protocol (Dvorak et al., 1988) and Ampure bead-based size selection instead of SYBR-gold agarose gel-based size selection. The prepared library was quantified using qRT-PCR using known concentration standards and sequenced on two lanes of a NovaSeq 6000 SP flow-cell using 200 cycles (2 × 100 bp paired-end mode).

### Data processing and enrichment efficiency calculation

The adapter and low quality bases from raw RenSeq reads were removed using cutadapt v1.16 (Martin, 2011) with a minimum read length of 30 bp after trimming. The quality check for adapter and quality trimming was carried out using FastQC v0.11.7 (https://www.bioinformatics.babraham.ac.uk/projects/fastqc/). Trimmed reads were aligned against the reference genome assembly of cv. Chinese Spring (RefSeq v1.0, The International Wheat Genome Sequencing Consortium (IWGSC) (2018)) using BWA-MEM v0.7.17 (Li, 2013) with default parameters. The output was converted to binary alignment map (BAM) format using SAMtools v1.9 (Li et al., 2009) and then the sorting was carried out using NovoSort (V3.06.05). Sequences from the bait library were aligned to the RefSeq v1.0 using BLASTn v2.9.0 program (Altschul et al., 1990). Alignments with 95% identity and 70% query coverage were retained and alignments separated by 120 bp or less were merged using bedtools v2.29.2 (Quinlan and Hall, 2010). Finally, the total size of regions >= 100 bp in the reference genome covered by alignments to the baits was calculated and considered as the size of our capture target. The sorted BAM file of each genotype was used to calculate the number of reads mapped on target and on the whole genome using SAMtools. The enrichment factor (EF) was then determined as (N/M)/(T/G), where N is the number of reads mapped on target, M indicates the total number of mapped reads, T denotes the size of the targeted region and G is the size of the genome.

### De novo RenSeq assembly and NBS-LRR identification

Only genotypes with at least 1 million reads and an enrichment factor >=100 were considered. The quality trimmed data from each genotype was assembled *de novo* with CLC Assembly cell (https://digitalinsights.qiagen.com/products-overview/discovery-insights-portfolio/analysis-and-visualization/qiagen-clc-assembly-cell/) using the parameters -w=64 - p fb ss 200 900. The contigs from each genotype were annotated with AUGUSTUS v3.3.145 (Stanke et al., 2006) using wheat gene models as training datasets, and contigs harboring complete genes were identified. Amino acid (AA), coding sequence (CDS) and transcript sequence for each complete gene were extracted using getAnnoFasta.pl script from the AUGUSTUS package. AA sequences of gene models were used to predict protein domains using the pfam_scan.pl script from PfamScan (Chojnacki et al., 2017), which searches FASTA sequences against the Pfam HMM database (Mistry et al., 2020). The script was run with sequence e-value cutoff of 10^−5^ and domain e-value cutoff 0.2 keeping other parameters to default. Genes containing at least one NB-ARC (NBS) domain (pfam ID PF00931.23) were considered as NLRs and used for downstream analysis. There is no standard tool available to predict coiled coil (CC) domain, therefore all NLRs with the “Rx_N” (PF18052) domain which is predicted as coiled coil were classified as coiled coil (CC). The sequences were further classified as NBS (only NBS domain), NLs (NBS + LRRs), CNs (Rx_N + NBS), CNLs (Rx_N + NBS + LRRs), XN (Integrated domain (ID) + NBS), XNL (ID + NBS + LRR), XRN (ID + Rx + N), XCNL (ID + Rx_N + NBS + LRR) based on domain composition. A bash script was used to retrieve gene structure information such as gene length, number of exons and introns from GFF files. The CDS sequences of NLRs from each genotype were aligned against RefSeq v1.0 using GMAP (Wu and Watanabe, 2005). The alignments were filtered with 70% query coverage and 95% identity cutoff.

### Clustering and saturation analysis

The AA sequences of all the NLRs identified from all genotypes were clustered using the easy-linclust workflow from MMseqs2 software suite (Steinegger and Söding, 2017). The program was run with e-value 1e-15, --min-seq-id 0.95, --seq-id-mode 2 (longer sequences), --cluster-mode 2 (coverage of query), --kmer-per-seq 100, keeping other parameters to default). The saturation analysis was carried out to determine the number of genotypes needed to capture 95% OGs. This was done by making random selection of the genotypes and counting the number of OGs present in these selections. The analysis was carried out separately for all the genotypes, only plant genetic resources and only using elite lines. The process was repeated 100 times starting with two and ending with a maximum number of genotypes for each category.

### Identification of elite and PGR-specific OGs

The genotype information of NLRs from each OG (variable *OG*). was used to classify the OG as specific to elite lines, specific to plant genetic resources (PGR) or common (variable *Type*). Further, the resistant OGs were identified based on the following linear model:

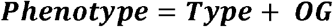

The OGs with negative effect and P value <0.01 were considered as resistant OGs. The number of resistant OGs from respective accessions were counted and correlation of OG count with disease susceptibility was studied.

### Chromosome scale genome assembly of “Attraktion” cultivar

The HiFi reads were assembled using hifiasm v0.14 (Cheng et al., 2021) to generate a primary contig assembly. The pseudomolecule construction was carried out using the TRITEX pipeline (Monat et al., 2019). For this, the guide map was constructed by aligning the single copy sequences from Julius to the Attraktion contig assembly. The Hi-C data was then used for chimera breaking and contig ordering to generate pseudomolecules. Transposable elements (TE) were annotated using a homology based approach implemented in RepeatMasker v4.0.8 (Smit et al., 2004). A custom library was created by downloading and combining wheat TE sequences from ClariTeRep: https://github.com/jdaron/CLARI-TE) and 2825 complete plant TE sequences (http://botserv2.uzh.ch/kelldata/trep-db/downloads/trep-db_complete_Rel-16.fasta.gz).

Gene annotation was carried out using AUGUSTUS v3.3.1. Initially, CDS sequences of high confidence (HC) genes from RefSeq v2.1 (Zhu et al., 2021a) were aligned to the Attraktion assembly using GMAP-GSNAP and Alignment was filtered with 70% coverage and 95% identity and top hits for each gene were extracted. The outputs of RepeatMasker and GMAP-GSNAP were combined and a GFF file was created. This GFF file along with wheat gene models served as a training dataset for AUGUSTUS.

The assembly completeness was assessed with 1,614 Benchmarking Universal Single Copy Orthologs (BUSCO v5.1.2) (Simao et al., 2015) genes from plants using “genome mode”. The assembly quality was also evaluated both as genome and gene level. For genome-wide comparison, single copy sequences from RefSeq v2.1 were aligned to Attraktion assembly using minimap2 v2.17 (Li, 2018). For gene level comparison, the transcript sequences of HC genes from RefSeq v2.1 were aligned against the transcript sequences of Attraktion using LAST (Kiełbasa et al., 2011). Alignment filtration and synteny analysis was carried out using MCScan (https://github.com/tanghaibao/jcvi/wiki/MCscan-%28Python-version%29).

The structural variations (SVs) were detected using the SyRI pipeline (Goel et al., 2019) with default parameters. For this, the Refseqv2.1 assembly was aligned to the Attraktion assembly using unimap, a fork of minimap2 optimized for assembly-to-reference comparison. The script “sam2delta.py” from RaGOO (Alonge et al., 2019) was used for SAM to mummer-delta format conversion. The delta file was filtered using delta-filter utility from MuMmer v4.0 (Marçais et al., 2018) to filter out smaller alignments (2000 bp) and the file was converted to TSV format using show-coords utility from MuMmer v4.0. The TSV file served as input for SyRI. The SVs from SyRI were reclassified into presence-absence variations (PAVs), inversions and translocations as follows: The CPL, DEL, DUP/INVDP (loss) variants, and the Attraktion sequences in NOTAL and TDM were converted as Absence SVs (relative to Attraktion). The CPG, INS, DUP/INVDP (gain) variants, and the query sequences in NOTAL and TDM were converted as Presence SVs (relative to Attraktion). The INV variants were regarded as inversions while the TRANS and INVTR were both regarded as translocation SVs.

### Phenotypic records and analyses

The experimental setup and quality assessment of yellow rust data were already presented in detail elsewhere (Schulthess et al., 2021). Briefly, five yellow rust artificially inoculated experiments plus two experiments relying on natural infections were conducted at five different German locations during harvest years 2019 and 2020. In three out of these seven field experiments, the presence of natural leaf rust infection was also recorded. Further details on these experiments can be found in **Table S3**. Yellow and leaf rust infection severity were expressed in a 1 (no symptoms) to 9 (severe infection) scoring scale according to the Protocols of the German Federal Plant Variety Office (http://www.bundessortenamt.de/internet30/fileadmin/Files/PDF/Richtlinie_LW2000.pdf). Outlier correction within and heritability within and across leaf rust experiments were assessed in the same way as for yellow rust (Schulthess et al., 2021). The best linear unbiased estimations (BLUEs) across experiments of yellow and leaf rust were used as phenotypes for downstream analyses (**Table S4**).

### Reference-based GWAS

The alignment records against the Chinese Spring reference in BAM format (see above) were used for variant identification. The variant calling was performed using the mpileup and call functions from SAMtools v1.9 and BCFtools (v1.8) (Li, 2011). The software was run with the - DV parameter for SAMtools mpileup and minimum read quality (-q) cutoff of 20. The bi-allelic SNPs were further filtered with minimum QUAL >=40; minimum read depth for homozygous call >=2 and minimum read depth for heterozygous calls >=4 using a custom AWK script. The commands were run in parallel wherever applicable to reduce computational time using GNU parallel (Tange, 2011).

The variant calling was also performed by aligning adapter and quality trimmed reads from each genotype against the genome assembly of Attraktion. Variant calling and SNP filtration were performed as described above except that minimap2 v.2.17 was used for mapping reads against the Attraktion assembly.

The GWAS for yellow rust and leaf rust was carried out using a univariate linear mixed model from GEMMA (v0.98) software (Zhou and Stephens, 2012) with -lmm 4 -miss 0.2 -maf 0.01 parameters, while keeping other parameters to default settings. Relatedness and population structure were accounted for using a kinship matrix in the form 2*(1-RD), where RD are the Rogers’ distances between genotypes computed from 17,840 high-quality GBS SNPs (Schulthess et al., 2021). GWAS was also carried out separately using PGRs and elite accessions to identify novel sources of resistance from PGRs, avoiding over-correction by the kinship matrix.

For paGWAS, SNPs with more than 20% missing data were scored as presence-absence variants. The presence-absence status of genotype calls was converted to reference/alternate allele calls. GWAS with these data were done with GEMMA as described above.

### Reference-free GWAS

The reference free GWAS was carried out using the kmersGWAS pipeline (Voichek and Weigel, 2020). Briefly, 31 bp *k*-mers that were supported by at least five reads were extracted using kmctools v3.1.1 (Kokot et al., 2017). The *k*-mers from all the genotypes were combined and a non-redundant *k*-mer presence-absence genotype matrix was generated. The pipeline was run with 100 permutations, the 5 million top *k*-mers and minor allele frequency 0.01, while setting other parameters to default. The significance threshold was determined by selecting 5th top P-value from the 100 top P values obtained from 100 permutations. To position the *k*-mers in the genome, they were mapped against the reference assemblies of Chinese Spring and Attraktion and positions of uniquely mapped *k*-mers were retrieved. The positional information along with p-values were used for generation of Manhattan plots using the qqman (Turner) R package.

The significant *k*-mers were also aligned to the NBS-LRR transcript database generated above and the number of *k*-mers aligned with 100% identity and 100% coverage to representative transcript sequences of each cluster were counted. The results were manually inspected and the candidate clusters with large proportions of significant *k*-mers for yellow rust and leaf rust were identified.

### Candidate gene identification

The results from various methods mentioned above were compared and consensus regions for yellow rust resistance were determined. The MCscan (https://github.com/tanghaibao/jcvi/wiki/MCscan-(Python-version)) software with default parameters was used to study local synteny between different wheat genome assemblies. The CDS sequences of candidate clusters identified based on the kmerGWAS method were aligned separately to each of the consensus regions identified for yellow rust using GMAP (Wu and Watanabe, 2005) and candidate genes were identified. The AA sequences of the NBS domain of candidate genes were extracted and aligned with NBS domains of cloned R genes from wheat using MAFFT v7.305 (Katoh et al., 2002). The spurious sequences or poorly aligned regions were removed using trimAl v1.2 (Capella-Gutiérrez et al., 2009). The phylogenetic analysis was carried out using IQ-TREE v1.6.12 (Nguyen et al., 2015). The phylogenetic tree was visualized using ggtree (Yu et al., 2017).

## Data availability

RenSeq raw data is available from the European Nucleotide Archive (ENA) under accession PRJEB48219. BioSamples IDs of sequence read sets of each genotype are given in **Table S1**. Yellow and leaf rust phenotypes used for association scans are given in **Table S4**. The Attraktion genome assembly and the underlying raw data are available under ENA accession PRJEB48529. RenSeq assemblies and annotation are available from the Plant Genomics &Phenomics Research Data Repository (Arend et al., 2016) under DOI https://doi.ipk-gatersleben.de/DOI/bf84cc9a-fbd5-415e-8da9-dbad84aa7758/ca6637e0-8b28-4c38-a3be-fcf4aca24d44/2/1847940088. The DOI was registered with e!DAL (Arend et al., 2014).

## Acknowledgments

Our research was supported by a grant from the German Federal Ministry of Education and Research (BMBF) in frame of the project GeneBank2.0 (grant nos. FKZ031B0184B and FKZ031B0184A). We thank Susanne König, Jacqueline Pohl, Ines Walde, Manuela Knauft, Mary Ziems, Christoph Martin, Jelena Perovic and Johannes Schneider for technical assistance. We are grateful to Anne Fiebig and Daniel Arend for their support in data submission.

## Author contributions

JCR, NS and MM designed research. AWS, PHB and JS produced phenotypic data. SMK and AH performed capture sequencing. AWS selected plant material and analyzed phenotypic data. SMK analyzed molecular data and performed association analyses. BS and BBHW provided the bait library. SP and AH generated PacBio HiFi and Hi-C data. SMK and MM constructed the reference assembly of cv. Attraktion. SMK, AWS and MM wrote the paper with input from all co-authors.

## Supplementary items

**Table S1**: Passport data and ENA accession numbers for 907 winter wheat accessions.

**Table S2**: RenSeq assembly statistics.

**Table S3**: Experimental setup and data quality assessment of leaf rust data.

**Table S4**: BLUEs of yellow and leaf rust used for association analyses.

**Figure S1**: **Distribution of NBS-LRR orthogroup size in the hexaploid wheat collection**. No orthogroups common to all the accessions were found because of random sequence drop out.

**Figure S2: Intrachromosomal Hi-C contact matrices for pseudomolecules of cv. Attraktion**.

**Figure S3: Whole-chromosome alignments of the pseudomolecules of cv. Attraktion to Chinese Spring RefSeq V2.1 assembly (Zhu et al. 2021)**.

**Figure S4**: **GWAS using SNPs identified relative to the reference assembly of Chinese Spring (RefSeq V1.0)**. GWAS was done in the panels: elite *(a)*, PGR *(b)*, and Elite and PGR combined *(c)*. No significant marker-trait associations were detected in the PGR panel at the 6A locus, indicating that the resistant haplotype is rare in PGR.

**Figure S5: Association scans for leaf rust resistance using different marker systems. (a)** SNPs identified relative to Chinese Spring RefSeq V1.0. *(b)* presence-absence GWAS using SNPs identified relative Chinese Spring RefSeq V1.0 and scored as presence-absence markers. *(c) k*-mers mapped against Chinese Spring RefSeq V1.0. Panels *(d), (e)* and *(f)* show the results of SNP-based, presence-absence and *k*-mer based GWAS when the reference sequence of cv. Attraktion was used for SNP identification or *k*-mer mapping. The blue horizontal lines indicate threshold above which associations are statistically significant.

## References

Alonge, M., Soyk, S., Ramakrishnan, S., Wang, X., Goodwin, S., Sedlazeck, F.J., Lippman, Z.B. and Schatz, M.C. (2019) RaGOO: fast and accurate reference-guided scaffolding of draft genomes. Genome Biol. 20, 224.

Altschul, S.F., Gish, W., Miller, W., Myers, E.W. and Lipman, D.J. (1990) Basic local alignment search tool. J Mol Biol 215, 403–410.

Arend, D., Junker, A., Scholz, U., Schüler, D., Wylie, J. and Lange, M. (2016) PGP repository: a plant phenomics and genomics data publication infrastructure. Database 2016.

Arend, D., Lange, M., Chen, J., Colmsee, C., Flemming, S., Hecht, D. and Scholz, U. (2014) e! DAL-a framework to store, share and publish research data. BMC bioinformatics 15, 214.

Arora, S., Steuernagel, B., Gaurav, K., Chandramohan, S., Long, Y., Matny, O., Johnson, R., Enk, J., Periyannan, S., Singh, N., Asyraf Md Hatta, M., Athiyannan, N., Cheema, J., Yu, G., Kangara, N., Ghosh, S., Szabo, L.J., Poland, J., Bariana, H., Jones, J.D.G., Bentley, A.R., Ayliffe, M., Olson, E., Xu, S.S., Steffenson, B.J., Lagudah, E. and Wulff, B.B.H. (2019) Resistance gene cloning from a wild crop relative by sequence capture and association genetics. Nat Biotechnol 37, 139–143.

Capella-Gutiérrez, S., Silla-Martínez, J.M. and Gabaldón, T. (2009) trimAl: a tool for automated alignment trimming in large-scale phylogenetic analyses. Bioinformatics 25, 1972–1973.

Cheng, H., Concepcion, G.T., Feng, X., Zhang, H. and Li, H. (2021) Haplotype-resolved de novo assembly using phased assembly graphs with hifiasm. Nature Methods 18, 170–175.

Chojnacki, S., Cowley, A., Lee, J., Foix, A. and Lopez, R. (2017) Programmatic access to bioinformatics tools from EMBL-EBI update: 2017. Nucleic Acids Research 45, W550–W553.

Della Coletta, R., Qiu, Y., Ou, S., Hufford, M.B. and Hirsch, C.N. (2021) How the pan-genome is changing crop genomics and improvement. Genome Biology 22, 3.

Dvorak, J., McGuire, P.E. and Cassidy, B. (1988) Apparent sources of the A genomes of wheats inferred from polymorphism in abundance and restriction fragment length of repeated nucleotide sequences. Genome 30, 680–689.

Gabur, I., Chawla, H.S., Liu, X., Kumar, V., Faure, S., von Tiedemann, A., Jestin, C., Dryzska, E., Volkmann, S., Breuer, F., Delourme, R., Snowdon, R. and Obermeier, C. (2018) Finding invisible quantitative trait loci with missing data. Plant Biotechnol J 16, 2102–2112.

Gaurav, K., Arora, S., Silva, P., Sánchez-Martín, J., Horsnell, R., Gao, L., Brar, G.S., Widrig, V., Raupp, J., Singh, N., Wu, S., Kale, S.M., Chinoy, C., Nicholson, P., Quiroz-Chávez, J., Simmonds, J., Hayta, S., Smedley, M.A., Harwood, W., Pearce, S., Gilbert, D., Kangara, N., Gardener, C., Forner-Martínez, M., Liu, J., Yu, G., Boden, S., Pascucci, A., Ghosh, S., Hafeez, A.N., O’Hara, T., Waites, J., Cheema, J., Steuernagel, B., Patpour, M., Justesen, A.F., Liu, S., Rudd, J.C., Avni, R., Sharon, A., Steiner, B., Kirana, R.P., Buerstmayr, H., Mehrabi, A.A., Nasyrova, F.Y., Chayut, N., Matny, O., Steffenson, B.J., Sandhu, N., Chhuneja, P., Lagudah, E., Elkot, A.F., Tyrrell, S., Bian, X., Davey, R.P., Simonsen, M., Schauser, L., Tiwari, V.K., Kutcher, H.R., Hucl, P., Li, A., Liu, D.-C., Mao, L., Xu, S., Brown-Guedira, G., Faris, J., Dvorak, J., Luo, M.-C., Krasileva, K., Lux, T., Artmeier, S., Mayer, K.F.X., Uauy, C., Mascher, M., Bentley, A.R., Keller, B., Poland, J. and Wulff, B.B.H. (2021) Evolution of the bread wheat D-subgenome and enriching it with diversity from <em>Aegilops tauschii</em>. bioRxiv, 2021.2001.2031.428788.

Goel, M., Sun, H., Jiao, W.-B. and Schneeberger, K. (2019) SyRI: finding genomic rearrangements and local sequence differences from whole-genome assemblies. Genome Biology 20, 277.

Hafeez, A.N., Arora, S., Ghosh, S., Gilbert, D., Bowden, R.L. and Wulff, B.B.H. (2021) Creation and judicious application of a wheat resistance gene atlas. Mol Plant 14, 1053–1070.

Hickey, G., Heller, D., Monlong, J., Sibbesen, J.A., Sirén, J., Eizenga, J., Dawson, E.T., Garrison, E., Novak, A.M. and Paten, B. (2020) Genotyping structural variants in pangenome graphs using the vg toolkit. Genome Biology 21, 35.

Hovmøller, M.S., Walter, S., Bayles, R.A., Hubbard, A., Flath, K., Sommerfeldt, N., Leconte, M., Czembor, P., Rodriguez-Algaba, J., Thach, T., Hansen, J.G., Lassen, P., Justesen, A.F., Ali, S. and de Vallavieille-Pope, C. (2016) Replacement of the European wheat yellow rust population by new races from the centre of diversity in the near-Himalayan region. Plant Pathology 65, 402–411.

Huang, L., Brooks, S.A., Li, W., Fellers, J.P., Trick, H.N. and Gill, B.S. (2003) Map-based cloning of leaf rust resistance gene Lr21 from the large and polyploid genome of bread wheat. Genetics 164, 655–664.

Jayakodi, M., Padmarasu, S., Haberer, G., Bonthala, V.S., Gundlach, H., Monat, C., Lux, T., Kamal, N., Lang, D., Himmelbach, A., Ens, J., Zhang, X.-Q., Angessa, T.T., Zhou, G., Tan, C., Hill, C., Wang, P., Schreiber, M., Boston, L.B., Plott, C., Jenkins, J., Guo, Y., Fiebig, A., Budak, H., Xu, D., Zhang, J., Wang, C., Grimwood, J., Schmutz, J., Guo, G., Zhang, G., Mochida, K., Hirayama, T., Sato, K., Chalmers, K.J., Langridge, P., Waugh, R., Pozniak, C.J., Scholz, U., Mayer, K.F.X., Spannagl, M., Li, C., Mascher, M. and Stein, N. (2020) The barley pan-genome reveals the hidden legacy of mutation breeding. Nature 588, 284–289.

Jupe, F., Witek, K., Verweij, W., Sliwka, J., Pritchard, L., Etherington, G.J., Maclean, D., Cock, P.J., Leggett, R.M., Bryan, G.J., Cardle, L., Hein, I. and Jones, J.D. (2013) Resistance gene enrichment sequencing (RenSeq) enables reannotation of the NB-LRR gene family from sequenced plant genomes and rapid mapping of resistance loci in segregating populations. Plant J 76, 530–544.

Katoh, K., Misawa, K., Kuma, K.-I. and Miyata, T. (2002) MAFFT: a novel method for rapid multiple sequence alignment based on fast Fourier transform. Nucleic Acids Res. 30, 3059–3066.

Kiełbasa, S.M., Wan, R., Sato, K., Horton, P. and Frith, M.C. (2011) Adaptive seeds tame genomic sequence comparison. Genome Res. 21, 487–493.

Kokot, M., Dlugosz, M. and Deorowicz, S. (2017) KMC 3: counting and manipulating k-mer statistics. Bioinformatics 33, 2759–2761.

Li, H. (2011) A statistical framework for SNP calling, mutation discovery, association mapping and population genetical parameter estimation from sequencing data. Bioinformatics 27, 2987–2993.

Li, H. (2013) Aligning sequence reads, clone sequences and assembly contigs with BWA-MEM. arXiv preprint 1303.3997.

Li, H. (2018) Minimap2: pairwise alignment for nucleotide sequences. Bioinformatics 34, 3094–3100.

Li, H., Feng, X. and Chu, C. (2020) The design and construction of reference pangenome graphs with minigraph. Genome Biology 21, 265.

Li, H., Handsaker, B., Wysoker, A., Fennell, T., Ruan, J., Homer, N., Marth, G., Abecasis, G. and Durbin, R. (2009) The sequence alignment/map format and SAMtools. Bioinformatics 25, 2078–2079.

Liu, F., Zhao, Y., Beier, S., Jiang, Y., Thorwarth, P., Cf, H.L., Ganal, M., Himmelbach, A., Reif, J.C. and Schulthess, A.W. (2020a) Exome association analysis sheds light onto leaf rust (Puccinia triticina) resistance genes currently used in wheat breeding (Triticum aestivum L.). Plant Biotechnol J 18, 1396–1408.

Liu, Y., Du, H., Li, P., Shen, Y., Peng, H., Liu, S., Zhou, G.-A., Zhang, H., Liu, Z., Shi, M., Huang, X., Li, Y., Zhang, M., Wang, Z., Zhu, B., Han, B., Liang, C. and Tian, Z. (2020b) Pan-Genome of Wild and Cultivated Soybeans. Cell 182, 162–176.e113.

Lopez-Arboleda, W.A., Reinert, S., Nordborg, M. and Korte, A. (2021) Global Genetic Heterogeneity in Adaptive Traits. Molecular Biology and Evolution.

López-Girona, E., Davy, M.W., Albert, N.W., Hilario, E., Smart, M.E.M., Kirk, C., Thomson, S.J. and Chagné, D. (2020) CRISPR-Cas9 enrichment and long read sequencing for fine mapping in plants. Plant Methods 16, 121.

Marçais, G., Delcher, A.L., Phillippy, A.M., Coston, R., Salzberg, S.L. and Zimin, A. (2018) MUMmer4: A fast and versatile genome alignment system. PLoS Comput. Biol. 14, e1005944.

Martin, M. (2011) Cutadapt removes adapter sequences from high-throughput sequencing reads. EMBnet. Journal 17, pp. 10–12.

Mascher, M., Wicker, T., Jenkins, J., Plott, C., Lux, T., Koh, C.S., Ens, J., Gundlach, H., Boston, L.B., Tulpová, Z., Holden, S., Hernández-Pinzón, I., Scholz, U., Mayer, K.F.X., Spannagl, M., Pozniak, C.J., Sharpe, A.G., Šimková, H., Moscou, M.J., Grimwood, J., Schmutz, J. and Stein, N. (2021) Long-read sequence assembly: a technical evaluation in barley. Plant Cell.

Michelmore, R.W. and Meyers, B.C. (1998) Clusters of resistance genes in plants evolve by divergent selection and a birth-and-death process. Genome Res 8, 1113–1130.

Mistry, J., Chuguransky, S., Williams, L., Qureshi, M., Salazar, Gustavo A., Sonnhammer, E.L.L., Tosatto, S.C.E., Paladin, L., Raj, S., Richardson, L.J., Finn, R.D. and Bateman, A. (2020) Pfam: The protein families database in 2021. Nucleic Acids Research 49, D412–D419.

Monat, C., Padmarasu, S., Lux, T., Wicker, T., Gundlach, H., Himmelbach, A., Ens, J., Li, C., Muehlbauer, G.J., Schulman, A.H., Waugh, R., Braumann, I., Pozniak, C., Scholz, U., Mayer, K.F.X., Spannagl, M., Stein, N. and Mascher, M. (2019) TRITEX: chromosome-scale sequence assembly of Triticeae genomes with open-source tools. Genome Biol 20, 284.

Nguyen, L.-T., Schmidt, H.A., von Haeseler, A. and Minh, B.Q. (2015) IQ-TREE: a fast and effective stochastic algorithm for estimating maximum-likelihood phylogenies. Mol. Biol. Evol. 32, 268–274.

Padmarasu, S., Himmelbach, A., Mascher, M. and Stein, N. (2019) In Situ Hi-C for Plants: An Improved Method to Detect Long-Range Chromatin Interactions. Methods Mol Biol 1933, 441–472.

Przewieslik-Allen, A.M., Wilkinson, P.A., Burridge, A.J., Winfield, M.O., Dai, X., Beaumont, M., King, J., Yang, C.-y., Griffiths, S., Wingen, L.U., Horsnell, R., Bentley, A.R., Shewry, P., Barker, G.L.A. and Edwards, K.J. (2021) The role of gene flow and chromosomal instability in shaping the bread wheat genome. Nature Plants 7, 172–183.

Quinlan, A.R. and Hall, I.M. (2010) BEDTools: a flexible suite of utilities for comparing genomic features. Bioinformatics 26, 841–842.

Sato, K., Abe, F., Mascher, M., Haberer, G., Gundlach, H., Spannagl, M., Shirasawa, K. and Isobe, S. (2021) Chromosome-scale genome assembly of the transformation-amenable common wheat cultivar ‘Fielder’. DNA Research 28.

Schulthess, A.W., Kale, S.M., Liu, F., Zhao, Y., Philipp, N., Rembe, M., Jiang, Y., Beukert, U., Serfling, A., Himmelbach, A., Fuchs, J., Oppermann, M., Weise, S., Boeven, P.H.G., Schacht, J., Longin, C.F.H., Kollers, S., Pfeiffer, N., Korzun, V., Lange, M., Scholz, U., Stein, N., Mascher, M. and Reif, J.C. (2021) GiPS: Genomics-informed parent selection uncovers the breeding value of wheat genetic resources. bioRxiv, 2021.2012.2015.472759.

Sears, E.R. and Miller, T.E. (1985) THE HISTORY OF CHINESE SPRING WHEAT. Cereal Research Communications 13, 261–263.

Simao, F.A., Waterhouse, R.M., Ioannidis, P., Kriventseva, E.V. and Zdobnov, E.M. (2015) BUSCO: assessing genome assembly and annotation completeness with single-copy orthologs. Bioinformatics 31, 3210–3212.

Smit, A., Hubley, R. and Green, P. (2004) RepeatMasker Open-4.0.

Stanke, M., Keller, O., Gunduz, I., Hayes, A., Waack, S. and Morgenstern, B. (2006) AUGUSTUS: ab initio prediction of alternative transcripts. Nucleic Acids Research 34, W435–W439.

Steinegger, M. and Söding, J. (2017) MMseqs2 enables sensitive protein sequence searching for the analysis of massive data sets. Nature Biotechnology 35, 1026–1028.

Steuernagel, B., Vrána, J., Karafiátová, M., Wulff, B.B.H. and Doležel, J. (2017) Rapid Gene Isolation Using MutChromSeq. In: Wheat Rust Diseases: Methods and Protocols (Periyannan, S. ed) pp. 231–243. New York, NY: Springer New York.

Tange, O. (2011) Gnu parallel-the command-line power tool. The USENIX Magazine 36, 42–47.

The International Wheat Genome Sequencing Consortium (IWGSC) (2018) Shifting the limits in wheat research and breeding using a fully annotated reference genome. Science 361, eaar7191.

Ting, J.P.Y., Lovering, R.C., Alnemri, E.S., Bertin, J., Boss, J.M., Davis, B.K., Flavell, R.A., Girardin, S.E., Godzik, A., Harton, J.A., Hoffman, H.M., Hugot, J.-P., Inohara, N., Mackenzie, A., Maltais, L.J., Nunez, G., Ogura, Y., Otten, L.A., Philpott, D., Reed, J.C., Reith, W., Schreiber, S., Steimle, V. and Ward, P.A. (2008) The NLR gene family: a standard nomenclature. Immunity 28, 285–287.

Turner, S.D. (2018) qqman: an R package for visualizing GWAS results using Q-Q and manhattan plots. The Journal of open source software.

Van de Weyer, A.L., Monteiro, F., Furzer, O.J., Nishimura, M.T., Cevik, V., Witek, K., Jones, J.D.G., Dangl, J.L., Weigel, D. and Bemm, F. (2019) A Species-Wide Inventory of NLR Genes and Alleles in Arabidopsis thaliana. Cell 178, 1260–1272 e1214.

Voichek, Y. and Weigel, D. (2020) Identifying genetic variants underlying phenotypic variation in plants without complete genomes. Nat Genet 52, 534–540.

Walkowiak, S., Gao, L., Monat, C., Haberer, G., Kassa, M.T., Brinton, J., Ramirez-Gonzalez, R.H., Kolodziej, M.C., Delorean, E., Thambugala, D., Klymiuk, V., Byrns, B., Gundlach, H., Bandi, V., Siri, J.N., Nilsen, K., Aquino, C., Himmelbach, A., Copetti, D., Ban, T., Venturini, L., Bevan, M., Clavijo, B., Koo, D.-H., Ens, J., Wiebe, K., N’Daye, A., Fritz, A.K., Gutwin, C., Fiebig, A., Fosker, C., Fu, B.X., Accinelli, G.G., Gardner, K.A., Fradgley, N., Gutierrez-Gonzalez, J., Halstead-Nussloch, G., Hatakeyama, M., Koh, C.S., Deek, J., Costamagna, A.C., Fobert, P., Heavens, D., Kanamori, H., Kawaura, K., Kobayashi, F., Krasileva, K., Kuo, T., McKenzie, N., Murata, K., Nabeka, Y., Paape, T., Padmarasu, S., Percival-Alwyn, L., Kagale, S., Scholz, U., Sese, J., Juliana, P., Singh, R., Shimizu-Inatsugi, R., Swarbreck, D., Cockram, J., Budak, H., Tameshige, T., Tanaka, T., Tsuji, H., Wright, J., Wu, J., Steuernagel, B., Small, I., Cloutier, S., Keeble-GagnÃre, G., Muehlbauer, G., Tibbets, J., Nasuda, S., Melonek, J., Hucl, P.J., Sharpe, A., Clark, M., Legg, E., Bharti, A., Langridge, P., Hall, A., Uauy, C., Mascher, M., Krattinger, S.G., Handa, H., Shimizu, K.K., Distelfeld, A., Chalmers, K., Keller, B., Mayer, K.F.X., Poland, J., Stein, N., McCartney, C.A., Spannagl, M., Wicker, T. and Pozniak, C.J. (2020) Multiple wheat genomes reveal global variation in modern breeding. Nature accepted.

Wu, T.D. and Watanabe, C.K. (2005) GMAP: a genomic mapping and alignment program for mRNA and EST sequences. Bioinformatics 21, 1859–1875.

Yu, G., Smith, D.K., Zhu, H., Guan, Y. and Lam, T.T.y. (2017) ggtree : an r package for visualization and annotation of phylogenetic trees with their covariates and other associated data. Methods in Ecology and Evolution 8, 28–36.

Yu, J., Pressoir, G., Briggs, W.H., Vroh Bi, I., Yamasaki, M., Doebley, J.F., McMullen, M.D., Gaut, B.S., Nielsen, D.M., Holland, J.B., Kresovich, S. and Buckler, E.S. (2006) A unified mixed-model method for association mapping that accounts for multiple levels of relatedness. Nat Genet 38, 203–208.

Zhang, J., Hewitt, T.C., Boshoff, W.H.P., Dundas, I., Upadhyaya, N., Li, J., Patpour, M., Chandramohan, S., Pretorius, Z.A., Hovmøller, M., Schnippenkoetter, W., Park, R.F., Mago, R., Periyannan, S., Bhatt, D., Hoxha, S., Chakraborty, S., Luo, M., Dodds, P., Steuernagel, B., Wulff, B.B.H., Ayliffe, M., McIntosh, R.A., Zhang, P. and Lagudah, E.S. (2021) A recombined Sr26 and Sr61 disease resistance gene stack in wheat encodes unrelated NLR genes. Nat Commun 12, 3378.

Zhou, X. and Stephens, M. (2012) Genome-wide efficient mixed-model analysis for association studies. Nature Genetics 44, 821–824.

Zhu, T., Wang, L., Rimbert, H., Rodriguez, J.C., Deal, K.R., De Oliveira, R., Choulet, F., Keeble-Gagnère, G., Tibbits, J., Rogers, J., Eversole, K., Appels, R., Gu, Y.Q., Mascher, M., Dvorak, J. and Luo, M.-C. (2021a) Optical maps refine the bread wheat Triticum aestivum cv. Chinese Spring genome assembly. Plant J. 107, 303–314.

Zhu, T., Wang, L., Rimbert, H., Rodriguez, J.C., Deal, K.R., De Oliveira, R., Choulet, F., Keeble-Gagnère, G., Tibbits, J., Rogers, J., Eversole, K., Appels, R., Gu, Y.Q., Mascher, M., Dvorak, J. and Luo, M.C. (2021b) Optical maps refine the bread wheat Triticum aestivum cv. Chinese Spring genome assembly. Plant J 107, 303–314.

