## Supplementary Figures for "A catalogue of resistance gene homologs and a chromosome-scale reference sequence support resistance gene mapping in winter wheat"

### Slide 1
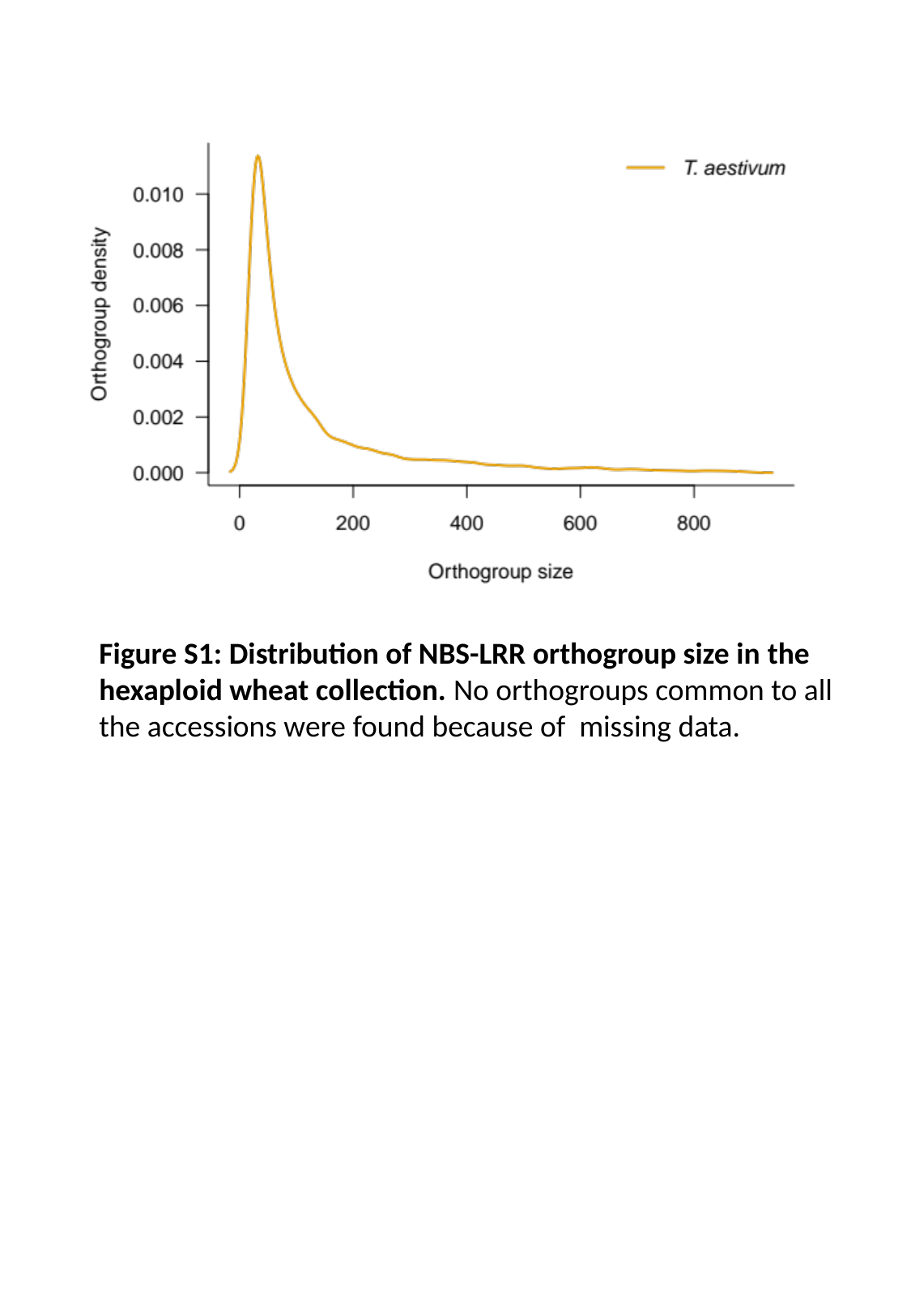

Figure S1: Distribution of NBS-LRR orthogroup size in the hexaploid wheat collection. No orthogroups common to all the accessions were found because of missing data.

### Slide 2
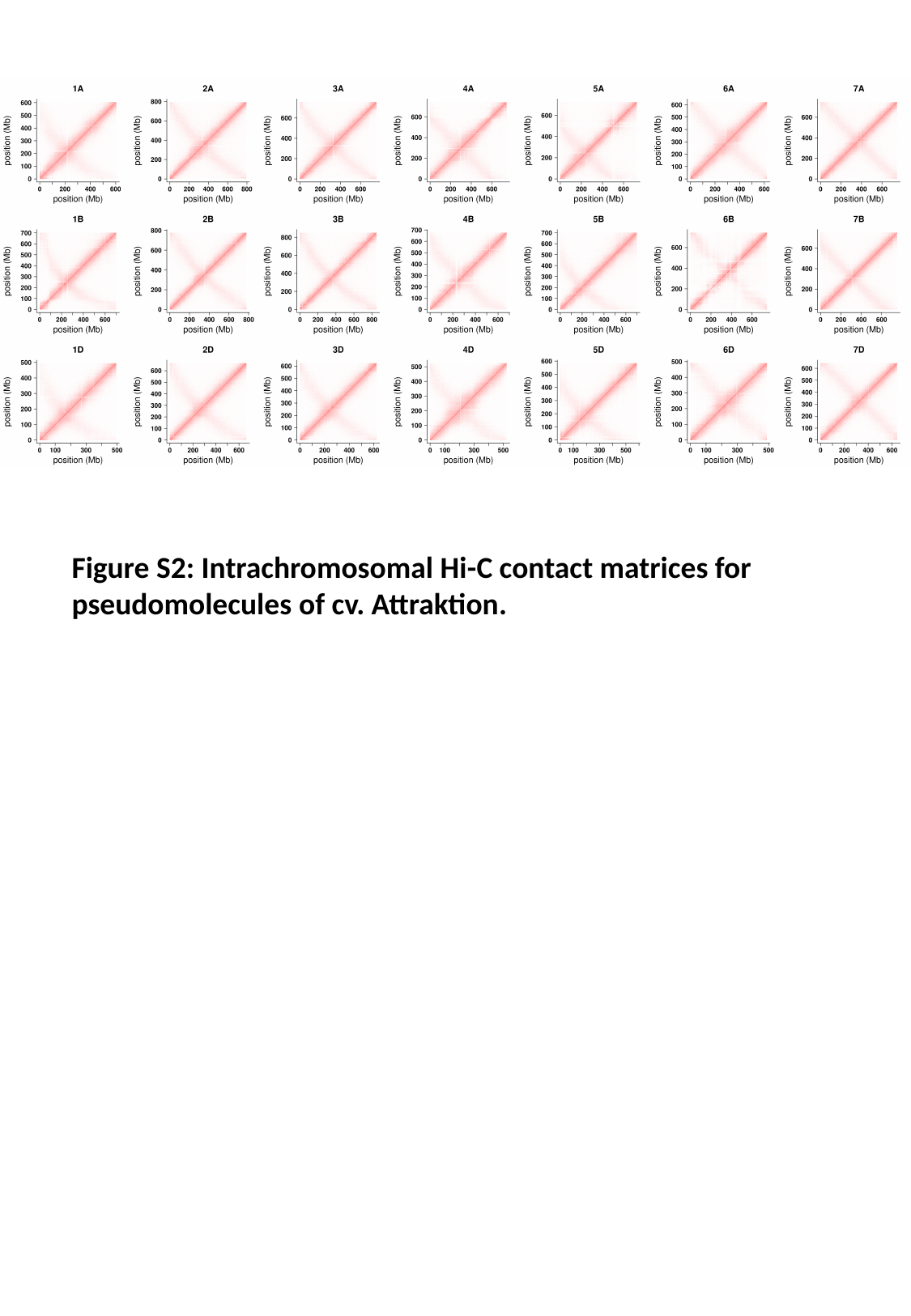

Figure S2: Intrachromosomal Hi-C contact matrices for pseudomolecules of cv. Attraktion.

### Slide 3
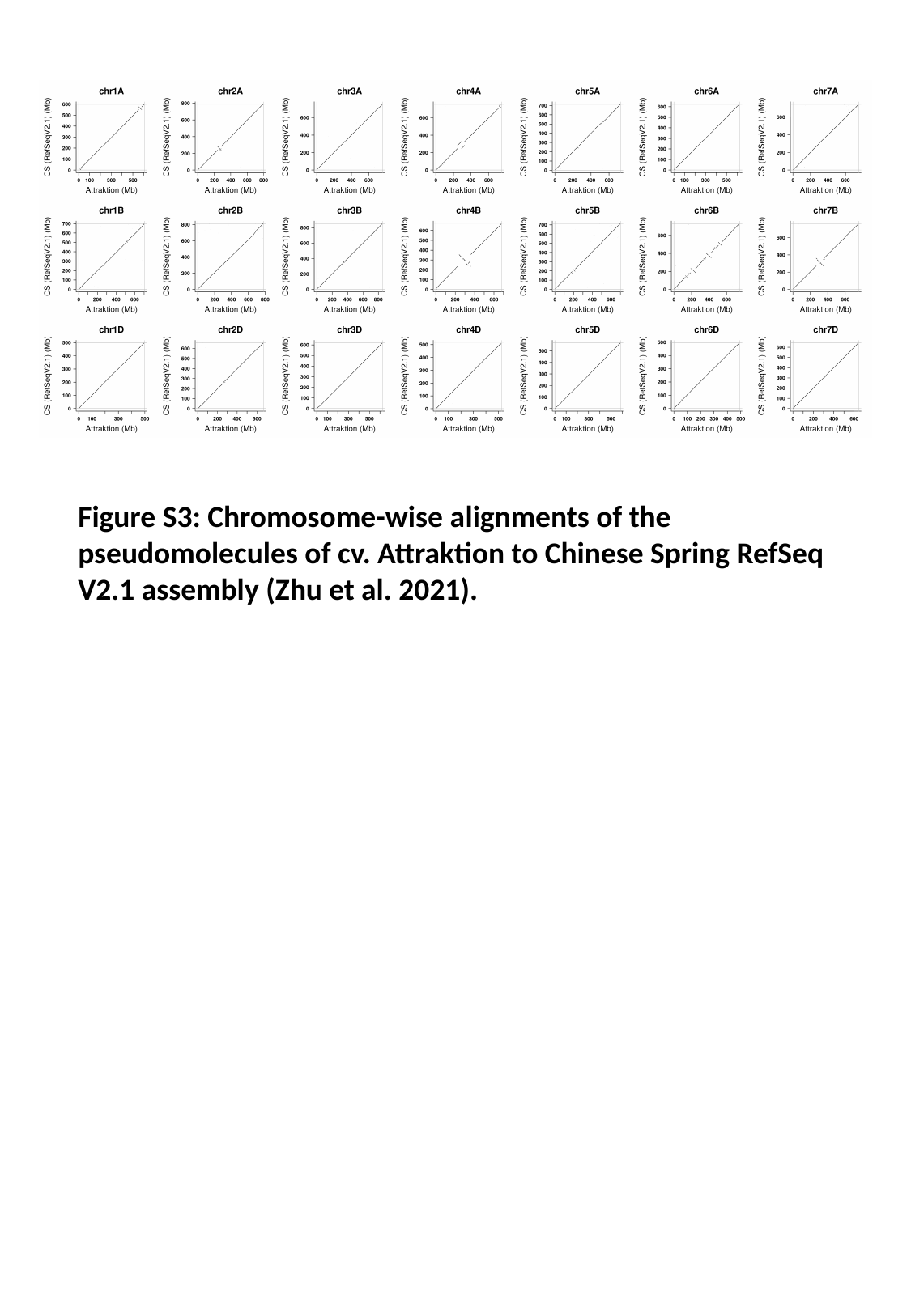

Figure S3: Chromosome-wise alignments of the pseudomolecules of cv. Attraktion to Chinese Spring RefSeq V2.1 assembly (Zhu et al. 2021).

### Slide 4
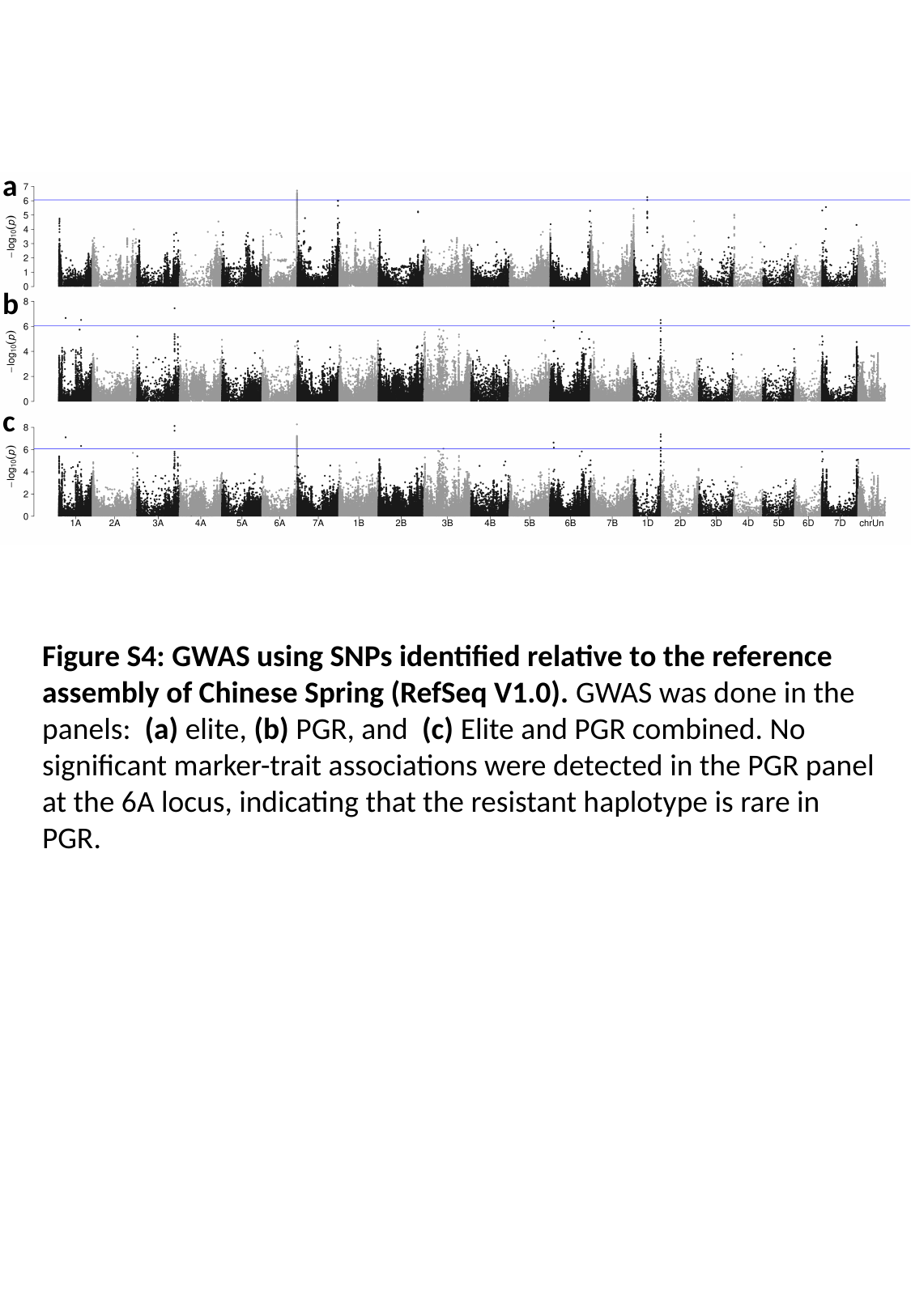

a
b
c
Figure S4: GWAS using SNPs identified relative to the reference assembly of Chinese Spring (RefSeq V1.0). GWAS was done in the panels: (a) elite, (b) PGR, and (c) Elite and PGR combined. No significant marker-trait associations were detected in the PGR panel at the 6A locus, indicating that the resistant haplotype is rare in PGR.

### Slide 5
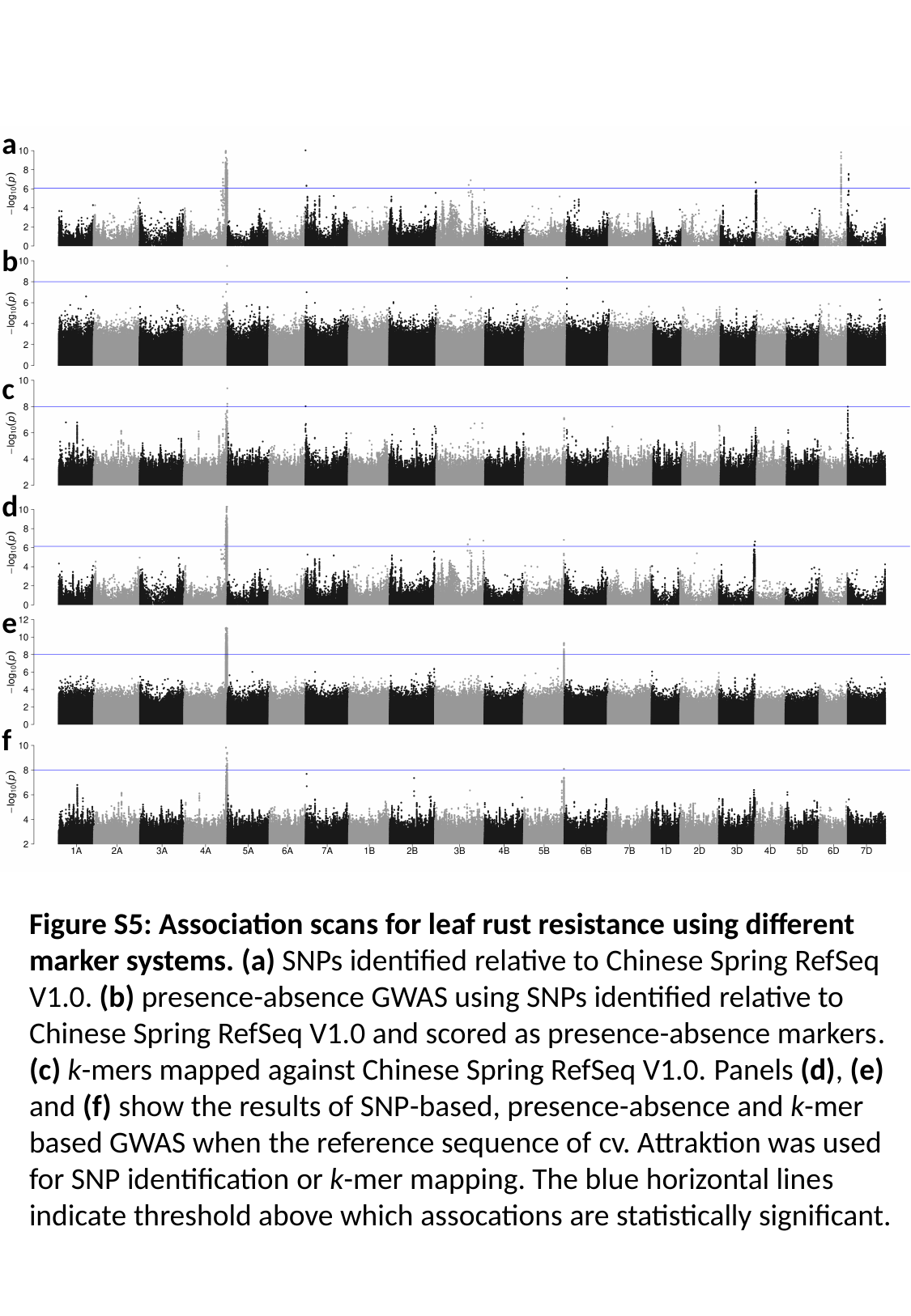

a
b
c
d
e
f
Figure S5: Association scans for leaf rust resistance using different marker systems. (a) SNPs identified relative to Chinese Spring RefSeq V1.0. (b) presence-absence GWAS using SNPs identified relative to Chinese Spring RefSeq V1.0 and scored as presence-absence markers. (c) k-mers mapped against Chinese Spring RefSeq V1.0. Panels (d), (e) and (f) show the results of SNP-based, presence-absence and k-mer based GWAS when the reference sequence of cv. Attraktion was used for SNP identification or k-mer mapping. The blue horizontal lines indicate threshold above which assocations are statistically significant.
